# Unsupervised learning for labeling global glomerulosclerosis

**DOI:** 10.1101/2024.09.01.610244

**Authors:** Hrafn Weishaupt, Justinas Besusparis, Cleo-Aron Weis, Stefan Porubsky, Arvydas Laurinavičius, Sabine Leh

## Abstract

Current deep learning models for classifying glomeruli in nephropathology are trained almost exclusively in a supervised manner, requiring expert-labeled images. Very little is known about the potential for unsupervised learning to overcome this bottleneck. To address this open question in a proof-of-concept, the project focused on the most fundamental classification task: globally sclerosed versus non-globally sclerosed glomeruli. The performance of clustering between the two classes was extensively studied across a variety of labeled datasets with diverse compositions and histological stains, and across the feature embeddings produced by 34 different pre-trained CNN models. As demonstrated by the study, clustering of globally and non-globally sclerosed glomeruli is generally highly feasible, yielding accuracies of over 95% in most datasets. Further work will be required to expand these experiments towards the clustering of additional glomerular lesion categories. We are convinced that these efforts (i) will open up opportunities for semi-automatic labeling approaches, thus alleviating the need for labor-intensive manual labeling, and (ii) illustrate that glomerular classification models can potentially be trained even in the absence of expert-derived class labels.

## 1. Introduction

With the advent of digital nephropathology, many diagnostic tasks in the evaluation of a kidney biopsy can now be addressed using artificial intelligence (AI), ushering in a new era of computer-assisted diagnosis [1]. For example, one of the most central tasks in the diagnosis of chronic kidney disease is the characterization of glomeruli, clusters of capillaries that act as the basic filtration units of the kidney and that can be affected by a myriad of disease-related morphological changes [2]. The classification of these structures, when done manually, can be quite time consuming and difficult, and has therefore been the focus of recent developments of deep learning (DL)-based nephropathology applications [3–8].

However, while clearly demonstrating the potential of DL for automatic classification of glomerular lesions, these works remain mostly proof-of-concept, lacking sufficiently large and diverse training datasets to achieve true clinical applicability. In fact, while whole slide scanners have enabled the rapid and feasible collection of vast amounts of histological image data, few of these data can be readily used to build DL models. Specifically, the training of such models is still mostly conducted in a supervised fashion, requiring additional image labels. However, histological images are inherently unlabeled, and the labeling is typically an extremely time-consuming task and highly dependent on domain expertise and thus remains a critical barrier to the development of AI tools.

Possible solutions for overcoming these bottlenecks include, for instance, the use of more interactive labeling strategies [9–12] or alternative “not-so-supervised” learning regimes [13], i.e. training strategies not explicitly relying on extensively labeled data. Out of such approaches, unsupervised and self-supervised learning approaches appear particularly appealing, because they can operate on completely unlabeled data and in the absence of supervision of a pathologist. Instead, utilizing a data-driven approach to finding a potentially meaningful separation of histological images, such approaches can either help to group images and patches [14–17] or the creation of pretrained/foundational models [18–20], which can be used as feature extractors or be repurposed for downstream tasks via transfer learning.

However, while a plethora of unsupervised and selfsupervised learning paradigms has been developed in the computer vision field [21–28], very little research has so far focused on the application of such approaches in the field of glomerular lesions. Liu et al. [29] employed selfsupervised learning on glomerular images, but only for the downstream classification between patches with and without glomeruli. Yao et al. [30] utilized self-supervised learning to pretrain a CNN on web-mined images of glomeruli, but subsequently still followed with a supervised fine-tuning using thousands of labeled images. Sato et al. [15] asked the question of whether a purely unsupervised definition of glomerular classes might provide a clinically meaningful basis for classifying glomerular changes in the absence of labels. While the study produced some promising concepts and initial results pertaining to the clustering of glomerular images, the separation between morphological classes and the purity of clusters appeared only limited and was not fully evaluated. Furthermore, the study utilized only a very restricted dataset, including only a single histological stain (Hematoxylin & Eosin; HE), and investigated only a single convolutional neural network (CNN) for feature extraction. Consequently, there is still a substantial lack of understanding regarding the full potential of clustering for the unsupervised distinction between glomerular lesions.

In theory, the clustering of morphological lesions might yield better results when performed on more extensive image datasets and/or when using a stain such as Periodic Acid Schiff (PAS) that better highlights relevant glomerular structures [31]. However, the majority of glomerular lesions might also be inherently difficult to distinguish via clustering, as the morphological changes in these cases are often segmental [32, 33], i.e. affecting only part of the glomerulus, while the remaining structure might be normal or present with other lesions. Thus, an unsupervised separation might be most successful for glomeruli with global lesions, i.e. with a morphological change affecting the entire glomerulus.

Consequently, towards investigating the feasibility of clustering glomerular lesions, it would be prudent to demonstrate it first in the context of such a well delineated task, i.e. a separation of the most distinct lesion categories, enabling a systematic proof-of-concept evaluation of methodology and performance. Specifically, the current study hypothesized that the best separation would probably be achieved for globally sclerosed (GS) glomeruli, as they represent a global pattern without any normal structures. This assumption is aligned with the observation that it is likely the class most easily classified [3–5]. The notion is further supported by the study by Altini et al. [7], who illustrated several CNN-based feature embeddings of glomerular images, where the most substantial separation appeared to manifest between the GS and non-globally sclerosed (nonGS) classes. Similarly, another recent publication showed that the unsupervised clustering of kidney biopsy image patches could theoretically distinguish between patches containing (globally) sclerosed and patches containing other glomeruli [14].

Beyond serving as a proof-of-concept and foundation for ongoing clustering efforts attempting the resolution of a broader range of morphological lesions, the unsupervised separation of GS and nonGS would also provide an avenue for semi-automatic labeling opportunities enabling the more straightforward collection of DL training datasets for building glomerulosclerosis classifiers. However, while the previous studies have demonstrated the general feasibility of such an approach, to the best of our knowledge, there is virtually no research systematically evaluating the clustering of GS and nonGS, including (i) an investigation of the most suitable strategy for clustering GS images, and (ii) explicitly documenting the accuracy with which such an unsupervised strategy can distinguish globally sclerosed glomeruli.

Accordingly, the current study thoroughly investigated the hypothesized possibility of detecting global glomerulosclerosis through clustering, by (i) utilizing larger datasets from varied sources and across different histological stains, (ii) evaluating a large number of feature extraction methods, (iii) carefully measuring the separation of classes in the associated feature embeddings, and (iv) explicitly evaluating the performance of clustering in capturing the classes.

## 2. Material and Methods

The project was conducted in four major stages as outlined in figure 1. Utilizing various repositories, a large collection of glomerular image patches was compiled (Fig. 1A) and preprocessed (Fig. 1B) for downstream analyses. Subsequently, numerous CNN models were employed to extract image features from the glomerular image patches, and the class separation between GS and nonGS in the resulting feature embedding was evaluated (Fig. 1C). Finally, it was investigated how accurately the GS and nonGS classes could be captured by clustering of the images in the feature space (Fig. 1D). A more detailed account of the individual steps is given below and in the Supplemental Material.

**Fig. 1.**
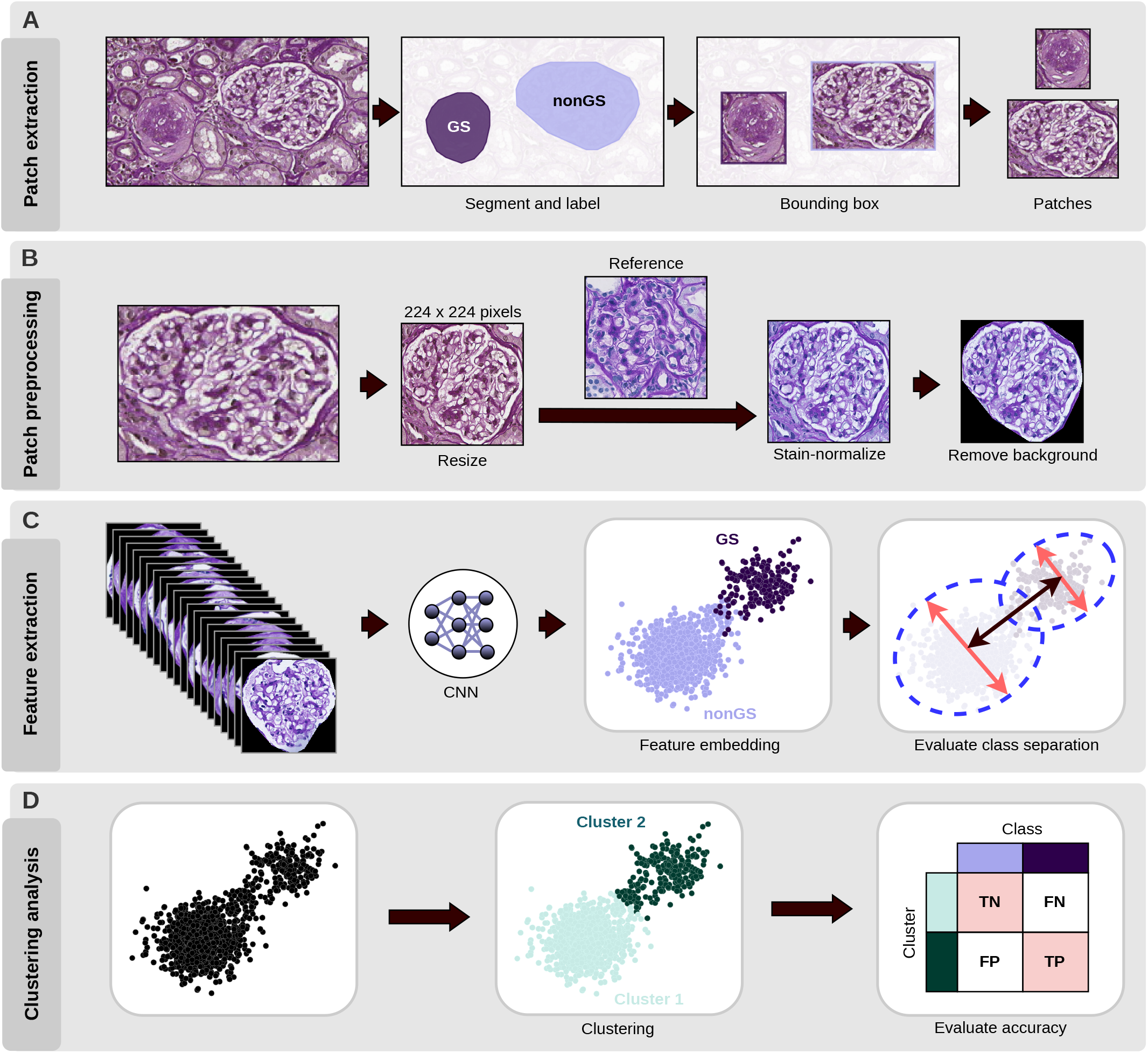
Workflow illustrating the overall procedure employed for data generation and analysis. **A**) Gathering glomerular patches from WSIs or other images: where necessary, glomeruli where segmented and labeled (here shown on part of a WSI from the Kidney Precision Medicine Project (KPMP) [34]); subsequently glomerular images where cropped using the bounding box. **B**) Glomerular image patch preprocessing: All image patches were first resized to 224 *×*224 pixels, then stain-normalized, and finally the surrounding tissue background was replaced by black pixels. **C**) Feature embedding: Each glomerular image patch was fed into a CNN for feature extraction, and in the resulting feature embedding for a single dataset, the separation between GS and nonGS images was evaluated quantitatively. **D**) Clustering of glomerular image patches: For each dataset, the images were clustered in the feature embedding, and the cluster assignments were compared to the corresponding class labels.

### 2.1 Data

#### Glomerular image datasets

The current study utilized glomerular image patches from seven sources: Besusparis et al. [5] (Besusparis2023, n=3993), Bueno et al. [35, 36] (Bueno2020, n=946), Gallego et al. [37] (two datasets: Gallego2021-HE/Gallego-PAS, n=611/527)[38], the Kidney Precision Medicine Project (KPMP, n=5978)[34], the Human BioMolecular Atlas Program (HuBMAP, n=4130)[39], the Norwegian Renal Registry (three datasets: NRR-PAS/NRR-HE/NRR-SIL, n=250/555/568), and Weis et al. [8] (Weis2022, n=5210). The sources displayed different properties, e.g. with respect to image formats, histological stains, availability of glomerular segmentations and/or class labels, which diagnoses and morphological lesions (beyond GS) are present in the dataset, and the proportion of GS and nonGS images, which are further described in the supplemental material and in supplementary tables 1 and 2.

#### Image patch generation and preprocessing

Depending on the source, the kidney biopsy images containing glomeruli came in different formats and were thus subjected to a sequence of preprocessing (Fig. 1A-B) to generate the final glomerular image patches.

Specifically, where not already available, glomeruli were segmented from the surrounding tissue and classified as either GS or nonGS (Fig. 1A). The segmentation annotations were utilized to compute the bounding box for each glomerulus, the image region within which was then extracted (Fig. 1A).

The resulting raw glomerular image patches were first rescaled to 224*×*224 pixels, and then, similar to the proce-dure documented by Sato et al. [15], image patches within each dataset were subjected to a stain-normalization step utilizing the Macenko method [40] (Fig. 1B). To evaluate the robustness of the feature embedding and clustering results with respect to the choice of reference image used during stain-normalization, 20 reference images with substantial color differences were selected per stain (Supp. Fig. 1) and utilized to generate 20 different stain-normalized versions of each dataset.

Finally, after image normalization, the surrounding tissue (pixels outside of the annotated glomerulus) in each image patch was masked out by replacing it with black color (Fig. 1B) [5, 7].

### 2.2 Feature extraction and evaluation

#### CNN models used for feature extraction

The current study compared the separation of GS and nonGS groups of glomeruli in the embedding of features obtained by different CNN models (Fig. 1C). A total of 34 CNNs were utilized (Supp. table 2), largely overlapping the models investigated for glomerular classification by Weis et al. [8] and Altini et al. [7], and also including the NASNetLarge model used in the previous clustering study by Sato et al. [15]. The CNN models were run either using the implementations available from Keras [41] or from the *classification_models* (https://github.com/qubvel/classification_models) library (Supp. table 3). All models were utilized without the top (classification) layers, the input tensor size was set to 224×224×3, weights pre-trained on ImageNet were used, and a global average pooling was applied to the output of the final convolutional layer.

#### Evaluating class separation in feature embedding

The separation of the nonGS and GS glomeruli in a given feature embedding was evaluated visually and using standard cluster evaluation metrics. Specifically, visual assessment involved the inspection of scatter plots after a Uniform Manifold Approximation and Projection (UMAP)[42] of the high-dimensional feature space down to two dimensions. For a quantitative assessment, the nonGS and GS labels were instead interpreted as cluster assignments, and then three internal cluster validity indices (CVIs) were applied to measure the quality of this “clustering” in the high-dimensional feature embedding: the Silhouette score [43], the C-index [44], and the Dunn index [45]. For the C-index, a better clustering would result in a lower score, while for the other two methods, a better clustering would result in a higher score.

### 2.3 Clustering analyses and evaluation

To cluster the glomerular images in the feature embedding (Fig. 1D), the project utilized either a Gaussian mixture model (GMM) strategy, similar to the study by [15], or a Leiden clustering [46] approach, in both cases aiming for exactly two clusters. The performance of clustering in capturing the nonGS and GS classes (Fig. 1D), was then evaluated (i) using confusion matrices, displaying the association between glomerular classes and the identified clusters, (ii) by one metric measuring the similarity between two clusterings, i.e. the adjusted Rand index (ARI) [47], and (iii) by one metric measuring classification performance, i.e. accuracy (ACC).

## 3. Results

### CNN-based feature embedding separates GS and nonGS

To evaluate the ability to automatically detect GS and nonGS glomerular images via unsupervised learning, the study first aimed to assess the separation of these classes in the feature space provided by different CNN models. Towards this end, ten datasets of glomerular images were selected, referred to respectively as Besusparis2023, Bueno2020, Gallego2021-HE, Gallego2021PAS, HuBMAP, KPMP-PAS, NRR-HE, NRR-PAS, NRR-SIL, and Weis2022, covering various histological stains and glomerular lesion categories (Supp. tables 1-2). Each dataset was labeled for GS and nonGS and was further stain-normalized with 20 different reference images (Supp. fig. 1), producing a total of 200 datasets for testing. Subsequently, 34 different CNNs (Supp. table 3) were utilized to extract image features from each of the datasets.

Evaluating the relationship between class affiliations and feature embeddings via internal CVIs, i.e. the Silhouette score [43], the C-index [44], and the Dunn index [45], the models displayed substantial differences in the separation achieved between the two classes (Fig. 2A-C, Supp. fig. 2). Specifically, the MobileNet, DenseNet169, DenseNet201, SE-ResNet101, and Xception were among the best performing models, while SE-ResNeXt50, EfficientNetV2M, EfficientNetV2L, VGG16, and VGG19 ranked generally the lowest.

**Fig. 2.**
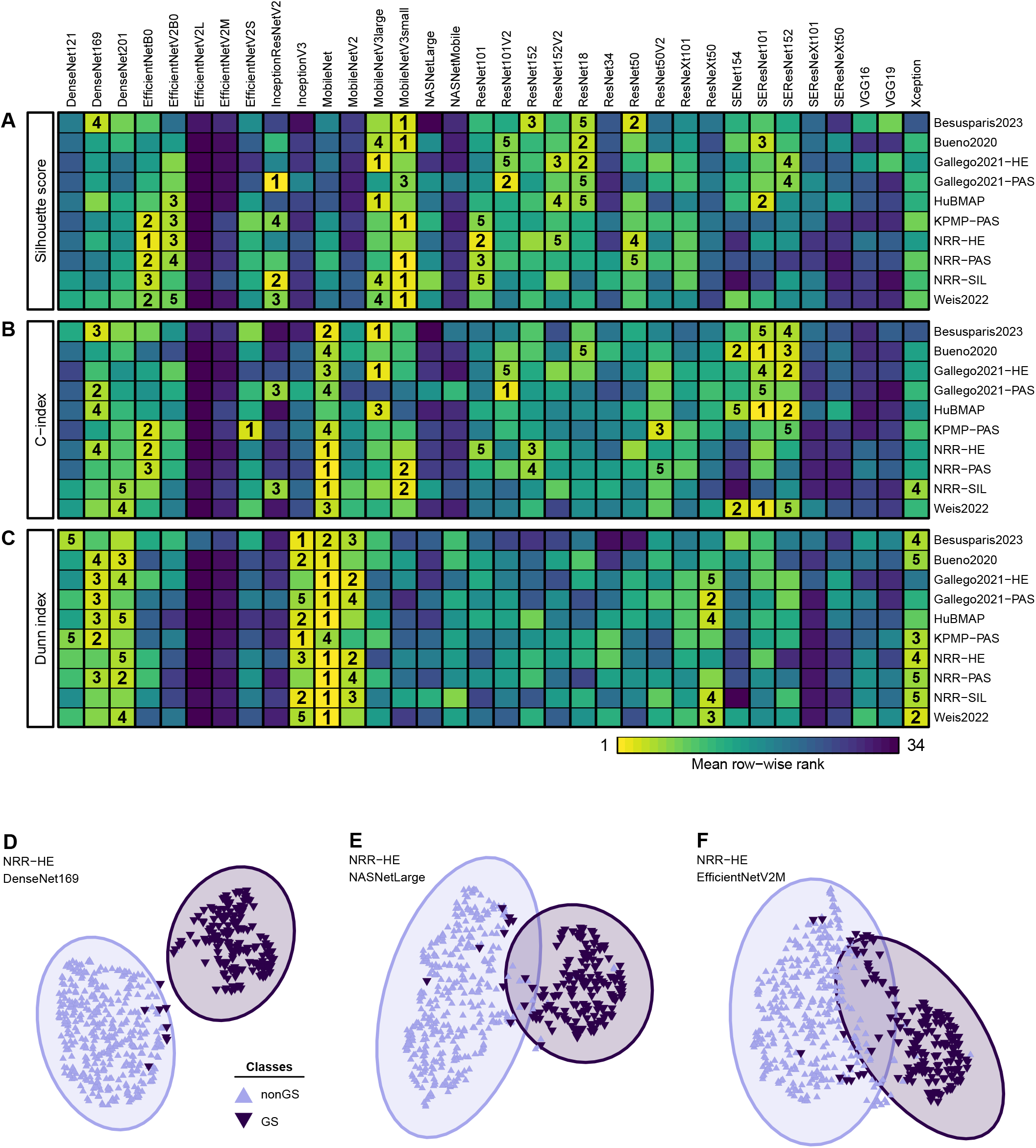
Evaluating the separation of GS and nonGS image patches between different CNN feature extractors. **A-C**) Heatmaps displaying the ranking of models based on internal cluster validity indices (CVIs), i.e. Silhouette score (A), C-index (B), and Dunn index (C). For each dataset, the models were ranked 20 times, once per stain-normalized version of the dataset, and the heatmap was generated based on the mean across these 20 rankings. **D-F**) UMAP embedding of the NRR-HE dataset using features from three different models, i.e. DenseNet169 (D), NASNetLarge (E), and EfficientNetV2M (F), chosen based on decreasing scores in all three CVIs. The shaded ellipses represent the 95% confidence ellipse for each class in the respective embedding.

When visually evaluated in respective twodimensional UMAP embeddings, models with different ranks according to the CVIs also displayed differences in the separation between GS and nonGS (Fig. 2D-F), suggesting that the ranking can provide an insight into how well the models could produce features separating between GS and nonGS. In addition, the normalization of datasets with different reference images also led to slight variations of the embedding (Supp. fig. 3).

Finally, picking two of the most promising feature extractors, i.e. the MobileNet and the DenseNet169, and visualizing the resulting UMAP embeddings, a marked separation between the GS and nonGS classes could be achieved across all datasets (Fig. 3, Supp. fig. 4), in comparison to the UMAP embeddings produced by some of the worst performing models such as the SE-ResNeXt50 or the EfficientNetV2L model (Supp. fig. 5-6).

**Fig. 3.**
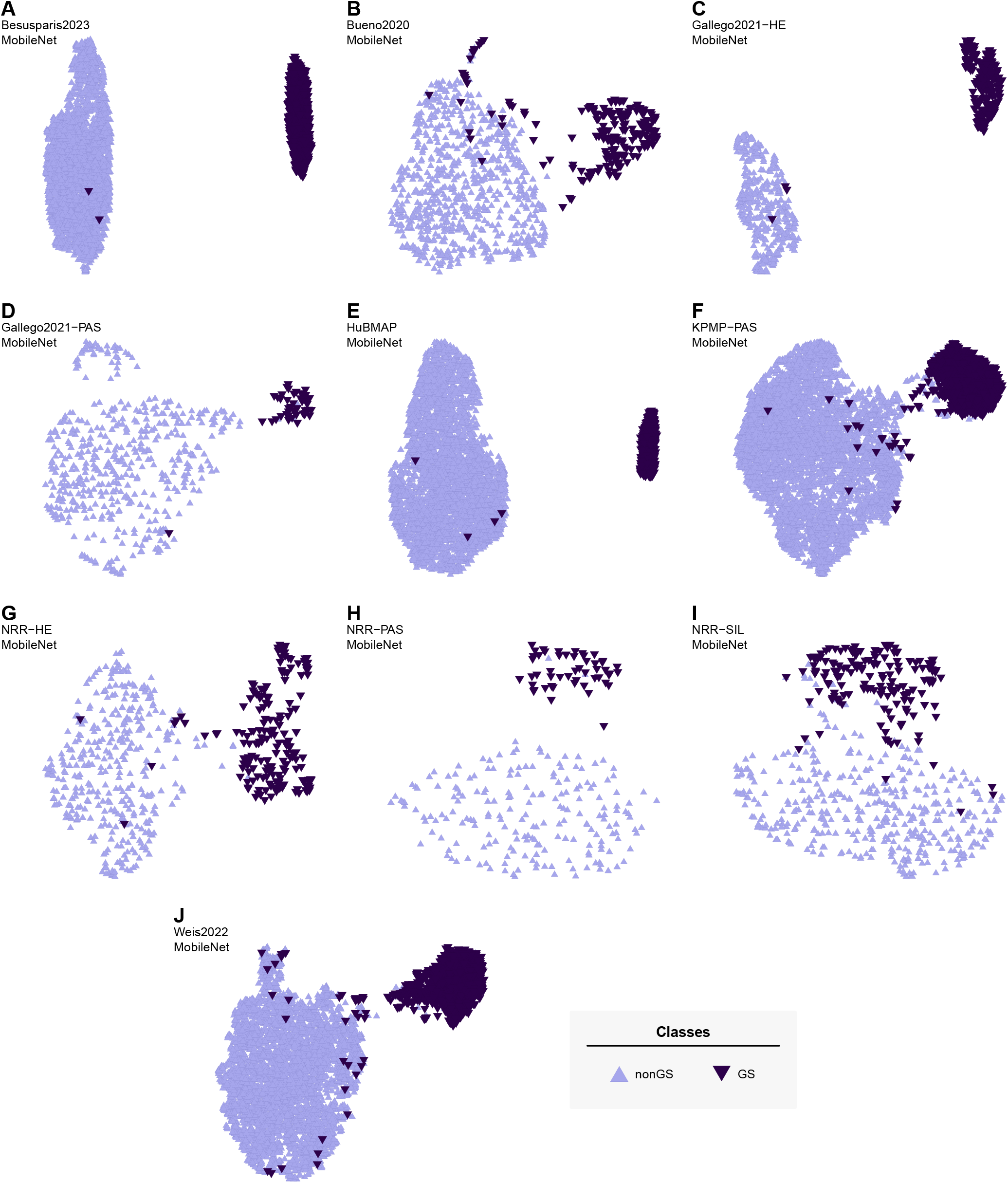
Scatterplots illustrating the separation of GS and nonGS glomerular image patches following feature extraction with the MobileNet and UMAP embedding in two dimensions. Each scatterplot depicts the feature embedding and classes of one stain-normalized variant of each of the ten datasets: Besusparis2023 (A), Bueno2020 (B), Gallego2021-HE (C), Gallego2021-PAS (D), HuBMAP (E), KPMP-PAS (F), NRR-HE (G), NRR-PAS (H), NRR-SIL (I), and Weis2022 (J). Light and dark purple colors indicate nonGS and GS glomerular images, respectively.

### Unsupervised learning captures GS and nonGS classes

Having demonstrated that CNN-derived features can enable a visually highly pronounced separation between the GS and nonGS classes, the next question was then whether a clustering strategy could also automatically detect the two glomerular classes from the respective feature embedding. Towards this goal, the project first adopted a strategy equivalent to what was proposed by Sato et al. [15], i.e. a UMAP projection followed by GMM clustering. Applied to the ten datasets, this approach generally produced good clustering results (data not shown). However, when investigating the robustness of this finding across the different stain-normalized versions of each dataset, there were cases in which GMM clustering failed to pick up the two visually apparent groups (Supp. fig. 7).

To overcome this issue, the project instead adopted a Leiden clustering strategy [46], which has recently also been shown highly feasible for the clustering of histological images in other studies [16]. Leiden clustering produced a similar separation of glomeruli (data not shown), but also managed to pick up the visually apparent groups of glomeruli in the cases in which GMM clustering failed (Supp. fig. 7, Supp. fig. 8). Consequently, Leiden clustering was then utilized as the method of choice for evaluating the clustering of GS and nonGS image patches.

Specifically, for each dataset, clustering was then performed on the features extracted by each of the 34 different CNNs, followed by the computation of the adjusted Rand index (ARI), comparing the ground truth labels (GS and nonGS) to the resulting cluster assignments (C1 and C2). Models were then ranked based on the observed ARI values (Fig. 4A), which suggested that the MobileNet might generally produce the best clustering performance across datasets. Particularly, assuming the minor cluster (C2) to capture the GS glomerular patches, and inspecting the corresponding confusion matrices for one stain-normalized version of each dataset, it was found that the use of the MobileNet in combination with Leiden cluster-ing led generally to few false-negative (FN, *≤*11%) and very few false-positive (FP, *≤*3.69%) detections (Fig. 4B-K). These findings were confirmed by inspecting the ARI (Fig. 4L) and balanced accuracy (Fig. 4M) values across all 20 stain-normalized versions for each dataset, with the mean balanced accuracies exceeding 93% in the Bueno2020 and NRR-SIL datasets, and exceeding 97% in all other datasets.

**Fig. 4.**
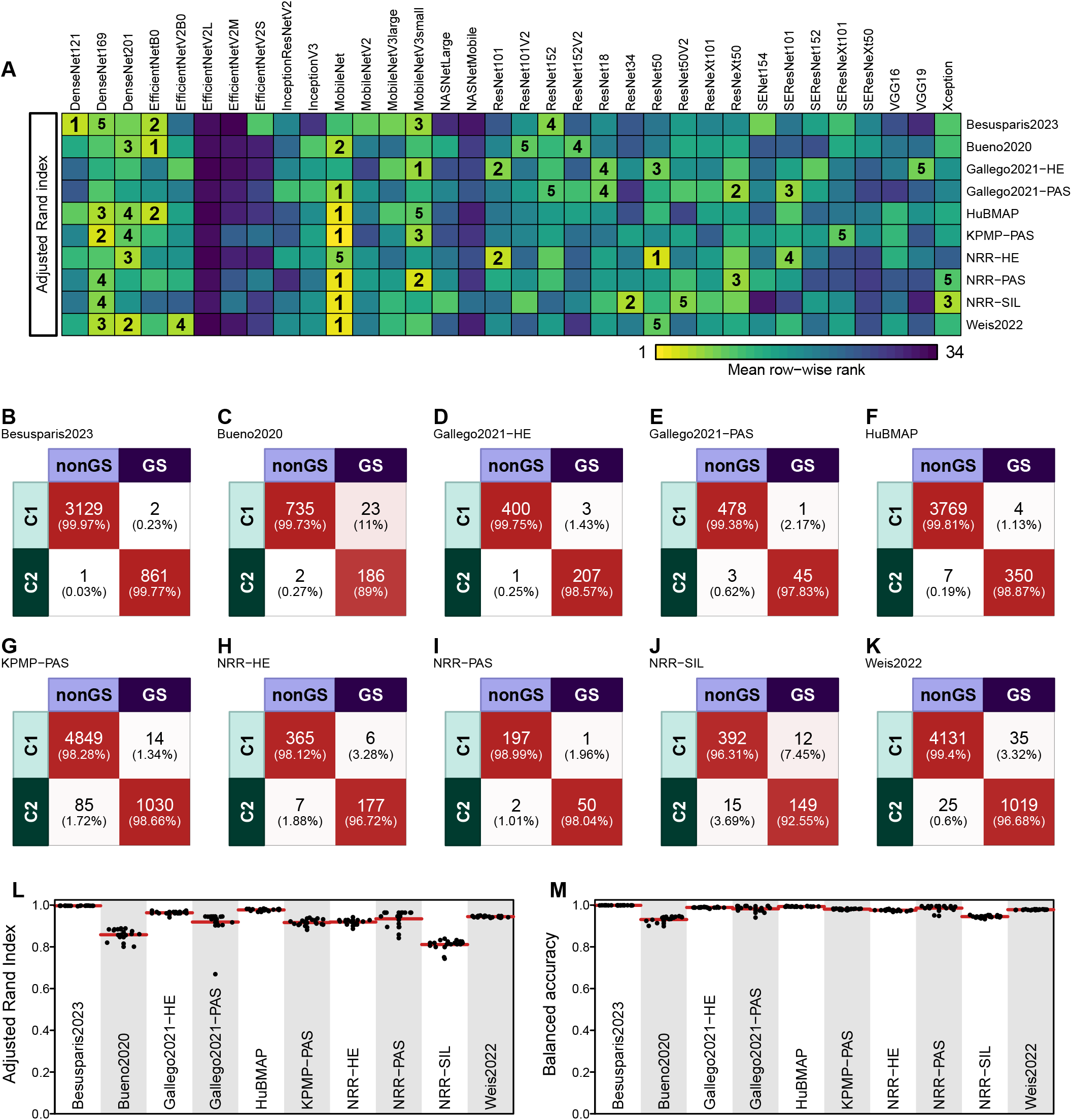
Evaluating the agreement between cluster and class labels. **A**) Heatmap displaying the ranking of CNN models within each dataset based on the adjusted Rand index (ARI). The ARI was computed between the class labels and the cluster assignments produced by Leiden clustering on the features produced by the respective CNN model. For each dataset, the models were ranked 20 times, once per stain-normalized version of the dataset, and the heatmap was generated based on the mean across these 20 rankings. **B-K**) Confusion matrices illustrating association between cluster (C1,C2) and class (nonGS,GS) memberships for the ten datasets: Besusparis2023 (B), Bueno2020 (C), Gallego2021-HE (D), Gallego2021-PAS (E), HubMap (F), KPMP-PAS (G), NRR-HE (H), NRR-PAS (I), NRR-SIL(J), and Weis2022 (K). **L-M**) Quantitative evaluation of the agreement between cluster affiliations and class labels as measured by the adjusted Rand index (ARI, L) and the balanced accuracy (ACC, M). Each data point represents the measurement performed on one of the 20 different stain-normalized variants of the respective dataset. The red lines indicate the mean across the 20 values for each dataset.

### Characterization and detection of misclustered cases

To understand the limitations of the clustering performances, a detailed investigation of the misclustered cases was conducted. Specifically, the analysis utilized a single stain-normalized version of each dataset, the MobileNet for feature extraction, and the Leiden algorithm for clustering. Subsequently, images labeled as GS and assigned to the major cluster (C1) where considered FNs, while images labels as nonGS and assigned to the minor cluster (C2) were considered FPs. A preliminary inspection of these images across all ten datasets (Supp. fig. 9-11) suggested that they constituted six different categories of glomerular images: (i) truly misclustered cases, (ii) mislabeled cases, (iii) borderline cases with advanced sclerosis (iv) globally sclerosed glomeruli with holes or split tissue, (v) images with potentially too little information to allow accurate labeling, e.g. tangential sections of glomeruli, and (vi) artifacts.

In addition, in the UMAP embedding, many of the misclustered images were also located on the border between the two clusters (Supp. fig. 9-11). We anticipated that these might predominantly be borderline cases either on the gradient between segmental and global sclerosis or otherwise difficult to distinguish from global sclerosis. Accordingly, we hypothesized that the cluster affiliation of these cases might not be robust across different clustering runs, which would enable their detection as uncertain cases. To investigate this question, we conducted two additional experiments, in which we compared either (i) the clustering of the 20 stain-normalized variants of the KPMP-PAS dataset after feature extraction with the MobileNet (Fig. 5A), or (ii) the clustering of one stain-normalized variant of the KPMP-PAS dataset processed with three different CNNs, i.e. DenseNet169, MobileNet, and MobileNetV3Small (Supp. fig. 12A). Uncertain cases could then be identified as those not always assigned to the same cluster across the different runs, and they covered many of the previously misclustered cases. Following a detailed re-labeling of these uncertain cases and the remaining FP and FN cases (those not absorbed into the uncertain group) by the consensus of two experienced nephropathologists, it was found that, in addition to distinct GS and nonGS images, they constituted many images with borderline GS, GS glomeruli with extensive white areas due to holes/split tissue, images with too little detail on the glomerulus for adequate labeling, or different types of artifacts (Fig. 5B). In addition, the re-labeling also suggested that many of the FN and FP clusterings were caused by a mislabeling in the intial labeling round (Fig. 5C-D, Supp. fig. 12B-C). The uncertain glomeruli were only very seldom mislabeled (Fig. 5E-F, Supp. fig. 12D-E). Glomeruli with more difficult appearances (borderline sclerosis, GS with holes, glomeruli with too little detail, or artifacts) were clearly present at varying percentages among all four groups of misclustered cases (Fig. 5C-F, Supp. fig. 12B-E), potentially explaining inconsistent clustering results for these cases.

**Fig. 5.**
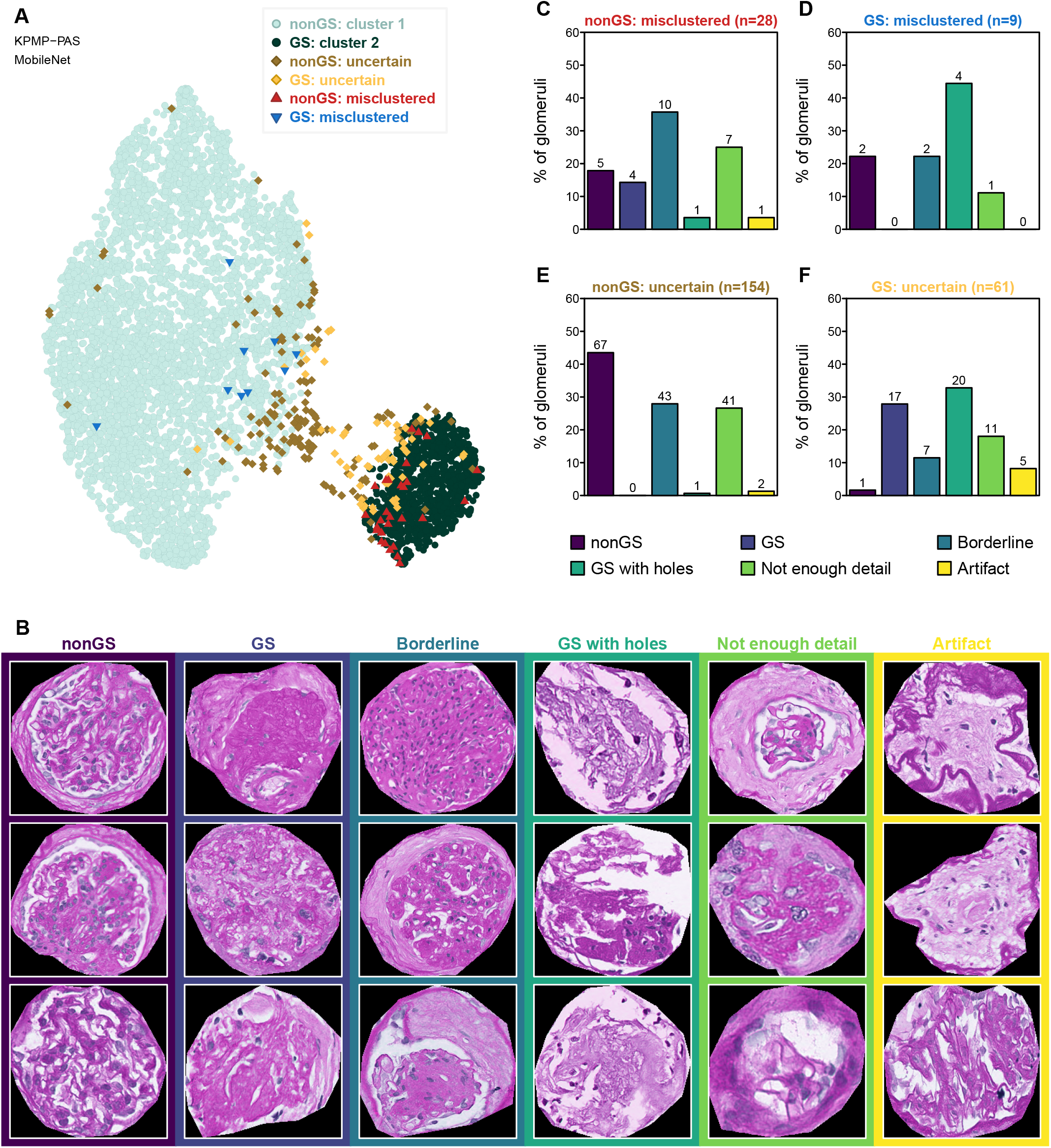
Evaluation of incorrectly clustered image patches: voting over multiple stain-normalized variations of the same dataset. **A)** Consensus of the clustering of the KPMP-PAS dataset following 20 different stain-normalizations and feature extraction using the MobileNet. Light and dark green circles indicate, respectively, glomeruli that are nonGS and always in cluster 1 and GS glomeruli that are always in cluster 2. Red and blue triangles indicate glomeruli that are always false-positives (nonGS in cluster 2) and false-negatives (GS in cluster 1), respectively. Brown and yellow diamonds indicate nonGS and GS glomeruli, respectively, that switch clusters between the different stain-normalizations and can thus be identified as uncertain cases. **B)** Three examples for each of the six different categories, i.e. nonGS, GS, borderline glomeruli with advanced sclerosis, GS with holes, glomeruli with insufficient detail for labeling, and artifacts, employed during relabeling of misclustered glomeruli. **C-F)** Barplots representing the percentages (absolute number above each bar) of these six groups among the glomeruli from each of the four misclustered categories, i.e. false-positives (C), false-negatives (D), and uncertain nonGS (E) and GS (F) glomeruli.

## 4. Discussions

The detection of global gomerulosclerosis has a clear role in CKD diagnostics, as it is one of the most prominent lesions inspected during biopsy reporting, and thus the automatic classification of GS glomeruli plays a central role in the development of computer-assisted diagnostic tools [4, 48]. However, current DL models for the classification of glomerular lesions are almost exclusively trained via supervised learning [3–8], which is reliant on extensively labeled image datasets, the establishment of which is often very costly. The current project demonstrated that, across several large collections of glomerular images, the GS and nonGS classes could be consistently separated using clustering, enabling the labeling of glomerular images through a purely data-driven, unsupervised approach.

Of note, the presented approach cannot be directly used in the diagnostic setting, as a single biopsy section typically contains too few glomeruli to warrant clustering. Instead, the outlined approach might help in easing the manual labeling requirements and/or enable downstream unsupervised learning strategies [15], pushing towards more robust and clinically relevant models for GS detection. Specifically, given the high accuracy with which GS images could be identified in the current study, the method appears highly suited for approaching a (semi-) automatic labeling of GS images, thus greatly simplifying the collection of DL training datasets.

The findings of the current study also lay the foundation for further studies, exploring the potential for unsupervised learning in detecting other, more challenging types of glomerular lesions. Specifically, automatic solutions for the labeling of morphological lesions beyond just GS and nonGS are in clear need in order to progress the training of clinically meaningful glomerular classification models. GS is likely the most pronounced and most easily classified lesion, and thus the results here serve only as the first proof-of-concept regarding the clustering of glomerular lesions. However, the accuracies achieved in the current study represents a promising starting point to investigate the further subclustering of both the GS, i.e. into obsolescent, solidified, and disappearing GS [30, 49], and the nonGS cluster into the various glomerular reaction patterns [2]. Achieving such a subclustering of meaningful nonGS lesion categories will likely require further efforts in identifying or establishing suitable feature extraction models able to capture features relevant to the separation of other morphological lesions.

Importantly, however, unsupervised approaches need to be approached with care. For instance, the current project always assumed all datasets to harbor a subset of globally sclerosed glomeruli, which were then assumed to constitute the minor cluster. In reality however, such a cluster-to-class assignment might fail if, for instance, no globally sclerosed glomeruli are present or are more abundant than the non-globally sclerosed glomeruli. Possible solutions might be to either inspect a few randomly sampled, representative cases from each cluster, or to include a set of external reference GS and nonGS images with known labels. Specifically, when processed together with the unlabeled dataset, including stain-normalization, feature extraction, and clustering, the cluster affiliation of the reference images might be used to automatically identify the GS and nonGS clusters. Alternatively, considering the high representational power of the CNNs like the MobileNet for clustering the classes, a few reference images might also be sufficient for fine-tuning such a CNN for a classification between GS and nonGS, and then predict labels on the unlabeled dataset directly.

Furthermore, the clustering of GS and nonGS might only work if a sufficient total number of glomeruli and sufficient number of GS image patches is included. The clustering might also be substantially affected by the composition of the dataset, including e.g. the types of diseases from which glomeruli were extracted, the types of glomerular lesions included, and the amount of pruning and curation performed on the image patches. Specifically, the best separations of the GS and nonGS classes were observed in the Besusparis2023 and HuBMAP datasets. While the former comprised a variety of different lesion categories, image patches were highly curated, being subjected to multiple rounds of validation to only include the image patches with the cleanest and most discernible lesions, thus potentially leading to the exceptional separation between GS and nonGS images. The HuBMAP data on the other hand appeared to lack, apart from globally sclerosed glomeruli, a large diversity of other lesions, thus leading to a clustering of almost exclusively GS and normal glomeruli, again resulting in a superb separability. The NRR, KPMP-PAS, Weis2022, and Bueno2020 datasets on the other hand were known or assumed to comprise both a broad range of diseases and/or lesion categories and were collected without much curation or pruning, thus representing a more natural distribution of glomerular appearances in both the GS and nonGS classes and consequently also a potentially more difficult task for clustering. In addition, the images in the Weis2022 dataset were also less clean with respect to glomerular extraction, containing sometimes only incompletely cropped glomeruli. The worst clustering performance was observed with respect to the NRR-SIL dataset. However, this finding was somewhat anticipated, seeing that the pathologist also had difficulties labeling some of the images purely based on an inspection of the Periodic Acid Schiff Methenamine Silver (SIL)-stained glomerulus itself, instead requiring a cross-check with corresponding PAS-stained images. Thus, if these images are inherently unsuited for accurate manual labeling by a pathologist, the clustering approach might experience similar difficulties in accurately separating these images. With respect to the evaluation of clustering performance, however, it should also be noted that the distinction between GS and segmental sclerosis (SS) might often be subjective. Specifically, the development of GS is a gradual process [50], and it might be difficult to consistently determine between what is still only SS and what is GS. Thus, manual labeling of these cases might be subject to inter-observer variability, which would also affect the ground truth and thus the evaluation of the clustering performance. In light of this consideration, a more complete evaluation of clustering performance might benefit from the labeling of glomeruli with any level of sclerosis, to be able to distinguish misclustered cases into true mistakes (e.g. nonsclerotic glomeruli in the GS cluster) and borderline cases. Importantly, the study also demonstrates that there is likely no one-fits-all CNN when it comes to feature embedding for clustering of GS and nonGS. Specifically, while the MobileNet appeared to be a generally good choice for feature extraction in combination with Leiden clustering, it was not the best choice across all datasets, with differences likely determined by variations in dataset composition, image properties, and histological stain. Thus, when applied to a new, unlabeled dataset, a strategy to find the feature extractor resulting in the best clustering might be to try different CNNs and evaluate the clustering quality via internal CVIs.

However, as indicated in the present study, selecting the best feature embedding and/or clustering based on internal CVI is not always straightforward. Specifically, there exists a vast number of internal CVI methods in the literature [51], each of which evaluates a specific objective function that might or might not align with the objective function optimized in the chosen clustering approach. In fact, in the current study, the ARI results for clustering with the Leiden algorithm did not correlate very well with the internal CVIs applied to the class separation in the feature embedding. Furthermore, CVIs might work differently depending on the skewness, noise, or subclusters in the data [51, 52]. For instance, in the context of determining the class separation between GS and nonGS, an embedding in which nonGS lesions are falling into discernible subclusters might result in a worse CVI as compared to an embedding with two compact clusters for GS and nonGS.

Finally, the current study was also subject to some technical and methodological limitations. Specifically, given the numerous combinations of models, datasets, and stain-normalizations utilized, the UMAP and GMM functions were run with default parameters to make them generically applicable rather than fine-tuning them separately for each individual case. In addition, only a single stain-normalization method was utilized. Considering the large number of alternatives approaches [53, 54], it would be worthwhile to investigate how the choice of stain-normalization methods affects the clustering of glomerular images. With respect to stain-normalization, the study also demonstrated that the choice of reference image can affect the outcome of the clustering. Thus, as shown here, a more robust labeling of GS might be achieved when incorporating a consensus over multiple stain-normalized variations of the dataset. While a set of diverse reference images in the current study was chosen solely based on a visual inspection, a more unbiased and automated way might be to pick them based on a computational evaluation of the stain style [55]. Similarly, it is not known how changes to the other preprocessing steps, i.e. the cropping and scaling or the removal of non-glomerular background, would influence clustering results. For instance, globally sclerosed glomeruli are often smaller than normal glomeruli [50], but by cropping and rescaling all glomeruli based on a bounding box rather than just using a fixed sized crop, information about glomerular sizes are lost. Since no golden standard for preprocessing of glomerular images prior to unsupervised learning appears available in the literature, it would be relevant to systematically evaluate the impact of alternative choices for each of these image preparation steps. Lastly, the study evaluated only the use of pre-trained CNN-based feature extractors, but did not explore other techniques for feature embedding such as autoencoders [15, 56] or other feature extractors trained via self-supervised learning on histological data [18–20]. Such models might be crucial to include in future investigations, when approaching the clustering of a more diverse set of glomerular lesion categories.

## 5. Conclusion

In summary, the current study clearly demonstrated that an unsupervised learning approach, utilizing CNNs pretrained solely on ImageNet, can separate globally and non-globally sclerosed glomerular images with very high proficiency. Specifically, among the evaluated CNN models, the MobileNet was found to generally perform best for clustering nonGS and GS glomeruli, achieving accuracies of over 95% in most datasets.

An inspection of the presumably misclustered glomeruli suggested that many such cases were either mislabeled during dataset generation or represented borderline glomeruli, inherently difficult to classify even by a pathologist. Furthermore, many such misclustered/uncertain glomeruli could be automatically detected by clustering the data multiple times, either using different stain-normalization references or different CNN networks for feature extraction. Following this approach, the accuracy of clustering could be increased even further, while highlighting the cases that likely require manual attention by a pathologist.

We are convinced that the outlined strategy will aid researchers (i) in achieving a more labeling-efficient generation of datasets for the training of deep-learning GS classifiers, and (ii) by providing a basis for ongoing research into the clustering of glomerular lesions. Specifically, considering the findings of the present work as a proof-of-concept, it appears promising to further explore the potential of unsupervised learning towards the detection of a broader range of glomerular lesions patterns.

## Acknowledgments

The results here are in part based upon data generated by (i) the Kidney Precision Medicine Project (https://www.kpmp.org), (ii) the HuBMAP program (https://hubmapconsortium.org), and (iii) the Norwegian Renal Registry (https://www.nephro.no/nnr.html). We would further like to thank Mindaugas Morku_nas for help in compiling the Besusparis2023 dataset. The project was funded by The Western Norway Health Authority (strategic research fund F-12563).

## Ethical declaration

Processing of NRR datasets: The collection and processing of data was approved by the Regional Committee for Medical and Health Research Ethics (REK number 517496).

Processing of Weis2022 dataset: The collection and processing of data was conducted in accordance with a vote from the ethics commission II of the Heidelberg university (vote 2020-847R).

Processing of Besusparis2023 dataset: The collection and processing of data was conducted with the permission from the Vilnius Regional Biomedical Research Ethics Committee (No. 2019/6-1148-637).

## Author contributions

*Study design:* H.W. *Data labeling and curation:* H.W., J.B., S.L. *Methodology, analysis, visualization:* H.W. *Writing:* H.W. *Manuscript review:* H.W., J.B., C.-A.W., S.P., A.L., S.L. *Data resources:* C.-A.W., S.P., A.L., S.L. *Supervision:* S.L.

