## Supplemental material for "Unsupervised learning for labeling global glomerulosclerosis"

**References** **20**

### 1. Material and methods

The following subsections provide an overview of the data and methods utilized during the study.

#### 1.1 Data

The project utilized various datasets of glomerular images collected from different resources, as further described below. Specifically, the utilized repositories comprised either readily available collections of glomerular patches, larger regions of kidney tissues including glomeruli, or kidney whole slide images (WSIs), from which glomeruli could be extracted through segmentation. An overview of the various datasets is given in supplementary table 1.

##### 1.1.1 Data sources

**1.1.1.1 Whole slide images** Raw kidney WSIs were obtained from three different sources, all with Periodic Acid Schiff (PAS) stain, including the Kidney Precision Medicine Project (KPMP, n = 218)[1], the Human BioMolecular Atlas Program (HuBMAP, n=14)[2], and the publication by Bueno et al. [3] (n=31, from here on referred to as Bueno2020 dataset). The KPMP-PAS and Bueno2020 WSIs comprised mostly needle biopsies with the occasional larger resection specimens (KPMP), while the HuBMAP WSIs comprised exclusively large resection specimens. Out of the 74 images available from HuBMAP, only the 14 images from formalin-fixed paraffin-embedded (FFPE) tissues were used. WSIs where available at resolutions of 0.2527 µm/pixel (40x magnification, KPMP), resolutions between 0.4936 and 0.4991 µm/pixel (20x magnification, Bueno2020), or at resolutions of 0.5000 µm/pixel (n=5, HuBMAP) or 0.6500 µm/pixel (n=9, HuBMAP).

**1.1.1.2 Pre-extracted glomerular images** Data from the other sources, i.e. Besusparis2023 [4], Gallego2021-HE & Gallego2021-PAS [5, 6], NRR-PAS & NRR-HE & NRR-SIL, and Weis2022 [7], were available as pre-extracted glomerular image patches. The Weis2022 dataset comprised rectangular patches including glomeruli; images were not always centered on or enclosing the entire glomerulus, and no segmentation masks were available. The NRR data comprised square image patches centered on glomeruli, also without accompanying segmentation mask. The Besusparis2023 dataset comprised square image patches centered on glomeruli and accompanied by a segmentation mask. The Gallego2021-HE and Gallego2021-PAS datasets comprised patches encompassing larger regions, typically including multiple glomeruli, and were accompanied by a segmentation mask for nonGS and GS glomeruli. The Weis2022, Gallego2021-PAS, and NRR-PAS datasets comprised images of PAS-stained glomeruli. The Gallego2021-HE and NRR-HE datasets contained images of Hematoxylin & Eosin (HE)-stained tissue. The NRR-SIL comprised images of Periodic Acid Schiff Methenamine Silver (SIL)-stained glomeruli. The Besusparis2023 dataset included images of glomeruli stained with a modified Picrosirius Red (mPSR).

##### 1.1.2 Glomerular segmentation and labeling

Some WSIs (HuBMAP) came with already established glomerular segmentations. For the other WSIs, glomeruli were either detected automatically through deep-learning-based segmentation (KPMP-PAS, Bueno2020) or segmented manually (HuBMAP). Finally, glomerular image patches from the Weis2022 and NRR-PAS/NRR-HE/NRR-SIL datasets were also obtained without accompanying glomerular segmentation annotations and where thus also segmented manually.

**1.1.2.1 DL-based glomerular segmentation** Glomerular candidates in the gathered collection of WSIs were automatically detected and segmented utilizing the Histo-Cloud web tool [8], using the *model-Glomeruli-11-13-20* model. For each WSI, the Histo-Cloud would provide instance segmentation results as a list of annotations, one per detected object and each in the form of a polygon defined by a set of border points.

KPMP-PAS WSIs were processed using default settings, i.e. using a downsampling factor of 2, while the Bueno2020 WSIs were processed using a downsampling factor of 1. Thus, with the original KPMP images coming at 40x magnification and Bueno2020 images coming at 20x magnification, both were segmented at 20x.

Importantly, glomeruli were automatically segmented from WSIs without any constraints/selections about regions of interests. Consequently, since WSIs often included more than one consecutively cut tissue section from the same biopsy, the resulting data set often contained multiple images for the same glomeruli. However, since none of the images were perfectly identical, none of the duplicates were removed.

All automatic segmentations were manually inspected and, if necessary, corrected in QuPath [9] by adjusting or replacing annotations with inaccurate boundaries, removing false-positive annotations, and adding annotations for missed glomeruli.

**1.1.2.2 Manual segmentation** For the HuBMAP dataset, seven WSIs had glomerular segmentations already available from an associated Kaggle challenge ("HubMAP - Hacking the Kidney")[10]. These WSIs were manually inspected in QuPath, and some missing glomeruli (mostly globally sclerosed glomeruli) were manually added as polygons. For the remaining seven HuBMAP WSIs, as well as the Weis2022 and NRR-PAS/NRR-HE/NRR-SIL datasets, glomeruli were also annotated manually in QuPath using the polygon tool.

**1.1.2.3 Labeling of glomeruli** The Weis2022 dataset comprised glomeruli already assigned to nine different classes, i.e. normal glomeruli, amyloidosis (AMY), GS, mesangial hypercellularity (MHC), membranoproliferative glomerulonephritis (MPGN), necrosis/crescent (NEC/CRE), nodular sclerosis (NS), not otherwise specified (NOS), and segmental sclerosis (SS). The NOS glomeruli were part of the original dataset collected in the study by Weis et al. [7], but were excluded during the experiments conducted in that paper.

A subset of the glomeruli from Besusparis et al. [4] was labeled into six classes, i.e. normal glomeruli, crescentic (CRE), GS, mesangioproliferative glomerulonephritis (MesPGN), MPGN, and SS. Specifically, the labeling was conducted as follows: (i) the glomerular lesion classifier developed in the publication was applied to the entire collection of pre-cropped glomerular images to predict class assignment probabilities, (ii) glomeruli with the highest predicted probabilities for each of the six classes were extracted, and (iii) the selected glomeruli were manually reviewed and labeled by an experienced nephropathologist (J.B.).

The NRR-PAS dataset included five classes of glomerular reaction patterns: AMY, GS, mesangial expansion (ME), NS of diabetic nephropathy, and SS. The NRR-HE dataset consisted of glomeruli labeled as: normal glomeruli, glomeruli with adherences (ADH), AMY, CRE, fibrinoid NEC, GS, ischemic changes (ISCH), ME, membranous glomerulonephritis (MGN), NS of diabetic nephropathy, proliferative changes, and SS. In this dataset, glomeruli could be assigned multiple labels. For example, glomeruli with crescents could also show fibrinoid necrosis. The “proliferative” category primarily included glomeruli with endocapillary hypercellularity (ECHC) but also encompassed those with a membranoproliferative reaction pattern. Additionally, some glomeruli with adherences were classified under segmental sclerosis. The NRR-SIL dataset was labeled as GS or nonGS, with these glomeruli retrieved from kidney biopsies of various non-neoplastic kidney diseases.

The datasets were intentionally selected because of or constructed to contain a more diverse set of morphological changes than just GS and normal glomeruli in order to emulate more clinically relevant testing conditions. However, for any downstream analyses, the labels from all of these datasets were then simplified into just the GS and nonGS classes.

For the remaining datasets (KPMP-PAS, HuBMAP, Bueno2020), glomerular images were labeled only as GS or nonGS.

**1.1.2.4 Validation of glomerular annotations and labels** The correction of automatic segmentation annotations, the manual segmentation, and the classification of glomeruli were conducted in two rounds. The initial segmentation and labeling of nonGS and GS was performed by the first author (non-pathologist) under the supervision of two experienced nephropathologists (S.L. and J.B.), after which all annotations and labels were double-checked and missing glomeruli annotated by an experienced nephropathologist (J.B.).

##### 1.1.3 Extraction and preprocessing of glomerular image patches

**1.1.3.1 Glomerular image patch generation** Glomerular image patches were obtained by first computing the bounding box for each glomerulus (based on the corresponding segmentation polygon). Subsequently, the image region included within this bounding box was extracted and resized to 224×224 pixels using cubic spline interpolation. Importantly, while some studies scale glomerular image patches while preserving the original aspect ratio [11], other studies appear to have scaled the image width and height independently without preserving aspect ratios [7, 12–14], which is also the approach employed in the current study.

Prior to extraction of glomerular patches from the larger Gallego2021 images, any glomeruli located directly on the border of the images were considered potentially incomplete and thus removed from the masks.

**1.1.3.2 Stain-normalization** After the generation of all image patches, a visual inspection revealed varying degrees of color variations among the images within each stain category (Supp. Fig. 1). Since variations in stain appearance might interfere with the clustering of the morphological classes, all glomerular image patches were stain-normalized via the Macenko method [15] implemented in the *tiatoolbox* (v1.4.0) package. Specifically, for each of the histological stains, 20 reference images were visually selected to cover a broad range of stain styles (Supp. Fig. 1A-D). Subsequently, each dataset was then normalized 20 times, once for each of the selected reference images for the respective stain, creating 20 different stain-normalized variations of the dataset.

#### 1.2 Analyses

To evaluate the ability of unsupervised learning to cluster GS and nonGS glomerular images, the current study was conducted in two main stages, i.e. (i) evaluating the separation between the two classes in the feature embedding obtained by a variety of CNN models, and (ii) investigating the agreement of the class labels with the cluster affiliations, obtained by clustering the images in the thus obtained feature embedding.

The project employed two different clustering strategies described in more detail below, i.e. (i) clusterings based on a Gaussian mixture model (GMM) approach, similar to the study by Sato et al. [16], and (ii) Leiden clustering [17].

##### 1.2.1 Evaluation of feature extraction

**1.2.1.1 Image feature extraction** DL-based image features were computed in Python via the Keras library with TensorFlow backend, using CNN models either implemented in Keras (https://keras.io/api/applications/), or the *classification_models* package (https://github.com/qubvel/classification_models). Models were configured to use a $224 \times 224 \times 3$ input tensor, use global average pooling after the final convolutional layer, and use the weights pre-trained on ImageNet.

For each dataset and model, prior to feature extraction, all images were first preprocessed using the model-specific *preprocess_input* function supplied with the respective model implementation.

**1.2.1.2 Evaluating class separation in feature embedding** The three internal cluster validity indices (CVIs), i.e. the Silhouette score [18], the C-index [19], and the Dunn index [20] were computed utilizing the implementation available from the *cluster_crit* (v1.0.1) library in python. CVIs were computed for each combination of one of the 20 stain-normalized datasets and one of the 34 CNN models used for feature extraction. For visualization in a heatmap, the results for a single CVI were ranked across all CNN models separately within each stain-normalized version of a dataset. Subsequently, within each dataset and CVI, the final score for a CNN was then obtained as the mean of the ranks observed across the 20 different stain variations of that dataset.

##### 1.2.2 Evaluation of clustering

**1.2.2.1 Dimensionality reduction** Data visualizations and GMM clustering were performed in a two-dimensional embedding of the high-dimensional feature vectors. The respective dimensionality reduction was performed via a two-step approach, first utilizing a principal component analysis (PCA) followed by a Uniform Manifold Approximation and Projection (UMAP)[21] step. The PCA was performed through the *PCA* and *fit_transform* functions (with default parameter settings) from the *scikit-learn* (v1.1.1) package, and UMAP was performed utilizing the *UMAP* and *fit_transform* functions (with default parameter settings) from the *umap-learn* (v0.5.3) package in Python. Confidence ellipses for the two groups of glomeruli in the UMAP projection were drawn using the *dataEllipse* function from the *car* (v3.1-2) library in R.

**1.2.2.2 Clustering of glomerular patches** GMM clustering was performed on the two-dimensional embedding of the feature vectors via the *GaussianMixture* function from the *scikit-klearn* (v1.1.1) package, setting only $n\_components = 2$, $n\_init = 100$, and $init\_params =$"$k - means + +$", while leaving all other parameters at default.

Leiden clustering was conducted utilizing functionalities from the *scanpy* (v1.9.5) package in python. Specifically, the raw feature vectors were first converted to an *AnnData* object, followed by the computation of a neighborhood graph (using default parameters). In order to cluster the neighborhood graph into two clusters, the Leiden clustering was then applied with incrementally increasing resolution values. Specifically, at very low resolution values, the algorithm would return only a single cluster, while increasing resolution values would lead to the discovery of an increasing number of clusters. For the datasets used in the current study, the selection of an adequate resolution value always resulted in the detection of two clusters, but the actual resolution value differed between glomerular dataset. Thus, an initial clustering was started at a sufficiently low resolution value, $r = 0.07$, and repeated for resolution values increased in steps of 0.005 until two clusters were returned. Apart from the resolution parameter, the clustering was performed using default parameters.

**1.2.2.3 Comparing cluster and class affiliations** For the comparison of cluster and class labels, the project used one metric measuring the similarity between two clusterings, i.e. the adjusted Rand Index (ARI) [22], and one metric measuring classification performance, i.e. accuracy (ACC). The ARI was computed utilizing the respective function from the R library *aricode* (v1.0.3). The ACC considered the clustering in the context of true positive and true negative predictions of global sclerosis, i.e. glomeruli in the minor cluster (C2) and labeled as GS were considered true positives (TPs) and glomeruli in the major cluster (C1) and labeled as nonGS were considered true negatives (TNs). The balanced ACC was then computed as:

$$ACC = \frac{\frac{|C1 \cap nonGS|}{|nonGS|} + \frac{|C2 \cap GS|}{|GS|}}{2} = \frac{\frac{|TN|}{|nonGS|} + \frac{|TP|}{|GS|}}{2}.$$

**1.2.2.4 Identification of inconsistently clustered glomerular images** To identify inconsistently clustered glomeruli, the current study inspected the robustness of cluster labels across repeated clusterings of a dataset by either (i) changing the stain-normalization reference utilized prior to feature extraction with the MobileNet, or (ii) using one stain-normalized version but extracting features with different CNN models. Specifically, images were considered to be consistently clustered if they received the same label in all clusterings across (i) the 20 stain-normalized versions of the dataset, or (ii) feature extractions obtained with three of the CNN models producing the best clustering results (DenseNet169, MobileNet, MobileNetV3Small), and were considered to be inconsistently clustered otherwise. The identification and subsequent analysis of such uncertain cases was conducted in the KPMP-PAS dataset, because it displayed the largest number of misclustered images.

#### 1.3 Figure generation

All figures displaying analysis results were generated in R (v.4.1.2).

### 2. Figures and tables

#### Supplementary table 1

**Supp. Table 1:** Properties of the various glomerular datasets

| Dataset | Ref. | Format* | Stain | Number of WSIs | Number of glomerular patches | Number of GS images | Segmented† | Classified§ | Classes‡ |
|---|---|---|---|---|---|---|---|---|---|
| **Besusparis2023** | [4] | Patches | mPSR | | 3993 | 863 | Before | Yes | Normal, CRE, GS, MesPGN, MPGN, SS |
| **Bueno2020** | [3, 24] | WSI | PAS | 31 | 946 | 209 | Histo-Cloud | No | |
| **Gallego2021-HE** | [5, 6] | Patches | HE | | 611 | 210 | Before | Yes | nonGS, GS |
| **Gallego2021-PAS** | [5, 6] | Patches | PAS | | 527 | 46 | Before | Yes | nonGS, GS |
| **HuBMAP** | [2] | WSI | PAS | 14 | 4130 | 354 | Before & Manually | No | |
| **KPMP-PAS** | [1] | WSI | PAS | | 5978 | 1044 | Histo-Cloud | No | |
| **NRR-PAS** | | Patches | PAS | | 250 | 51 | Manually | Yes | AMY, GS, ME, NS, SS |
| **NRR-HE** | | Patches | HE | | 555 | 183 | Manually | Yes | Normal, ADH, AMY, CRE, ECHC, GS, ISCH, ME, MGN, NEC, NS, SS |
| **NRR-SIL** | | Patches | SIL | | 568 | 161 | Manually | Yes | nonGS, GS |
| **Weis2022** | [7] | Patches | PAS | | 5210 | 1054 | Manually | Yes | Normal, AMY, GS, MHC, MPGN, NEC/CRE, NS, NOS, SS |

*: Images from the different sources were either provided in the form of WSIs or as pre-extracted image patches containing one or more glomeruli.

†: *Segmented* specifies whether the glomeruli were already segmented in the provided dataset (Before), were segmented automatically using the Histo-Cloud software [8] followed by manual correction (Histo-Cloud), or were segmented manually (Manually).

§: *Classified* Specifies whether the glomerular images were already classified in the provided dataset.

‡: ADH = glomeruli with adherences, AMY = amyloidosis, CRE = crescentic, ECHC = endocapillary hypercellularity, ISCH = ischemic, GS = global sclerosis, ME = mesangial expansion, MHC = mesangial hypercellularity, MesPGN = mesangioproliferative glomerulonephritis, MGN = membranous glomerulonephritis, MPGN = membranoproliferative glomerulonephritis, NEC = necrosis, NS = nodular sclerosis, NOS = not otherwise specified, SS = segmental sclerosis.

### Supplementary table 2

**Supp. Table 2:** Distribution of nonGS lesions in the datasets with available subclassifications

| Lesion category[†*] | Besusparis2023 | NRR-HE[§] | NRR-PAS | Weis2022 |
|---|---|---|---|---|
| **Normal** | 1530 | 146 | - | 1381 |
| **AMY** | - | 18 | 50 | 394 |
| **ADH** | - | 5 | - | - |
| **CRE** | 318 | 23 | - | - |
| **ECHC** | - | 28 | - | - |
| **ISCH** | - | 31 | - | - |
| **ME** | - | 91 | 49 | - |
| **MHC** | - | - | - | 946 |
| **MesPGN** | 727 | - | - | - |
| **MGN** | - | 24 | - | - |
| **MPGN** | 381 | - | - | 69 |
| **NEC** | - | 17 | - | - |
| **NEC/CRE** | - | - | - | 455 |
| **NS** | - | 1 | 50 | 233 |
| **NOS** | - | - | - | 174 |
| **SS** | 174 | 46 | 50 | 504 |

[†]: AMY = amyloidosis, ADH = adherence, CRE = crescentic, ECHC = endocapillary hypercellularity, GS = global sclerosis, ISCH = ischemic, ME = mesangial expansion, MHC = mesangial hypercellularity, MesPGN = mesangioproliferative glomerulonephritis, MGN = membranous glomerulonephritis, MPGN = membranoproliferative glomerulonephritis, NEC = necrosis, NS = nodular sclerosis, NOS = not otherwise specified, SS = segmental sclerosis.

[*]: The categories are based on the labels used in the four datasets. A blank field does not necessarily mean an absence of these glomeruli in the respective dataset, but the glomeruli with a more specific lesion might instead be included in a broader category, e.g. ME might include MesPGN cases, and the NEC/CRE includes both NEC and/or CRE cases.

[§]: The dataset included glomeruli with lesions from more than one category.

#### Supplementary table 3

**Supp. Table 3:** The CNN models used for feature extraction in the current study

| Implementation | Model name |
|---|---|
| **Keras**† | Xception |
| | VGG16 |
| | VGG19 |
| | ResNet50 |
| | ResNet50V2 |
| | ResNet101 |
| | ResNet101V2 |
| | ResNet152 |
| | ResNet152V2 |
| | InceptionV3 |
| | InceptionResNetV2 |
| | MobileNet |
| | MobileNetV2 |
| | DenseNet121 |
| | DenseNet169 |
| | DenseNet201 |
| | NASNetMobile |
| | NASNetLarge |
| | EfficientNetB0 |
| | EfficientNetV2B0 |
| | EfficientNetV2S |
| | EfficientNetV2M |
| | EfficientNetV2L |
| ***classification_models***§ | ResNet18 |
| | ResNet34 |
| | ResNeXt50 |
| | ResNeXt101 |
| | SE-ResNet18 |
| | SE-ResNet34 |
| | SE-ResNet50 |
| | SE-ResNet101 |
| | SE-ResNet152 |
| | SE-ResNeXt50 |
| | SE-ResNeXt101 |
| | SENet154 |

†: Utilizing the models and *preprocess_input* functions implemented in Keras: https://keras.io/api/applications/, last accessed 2024-03-24.

§: Utilizing the models and *preprocess_input* functions implemented in the *classification_models* package: https://github.com/qubvel/classification_models, last accessed 2024-03-24.

#### Supplementary figure 1

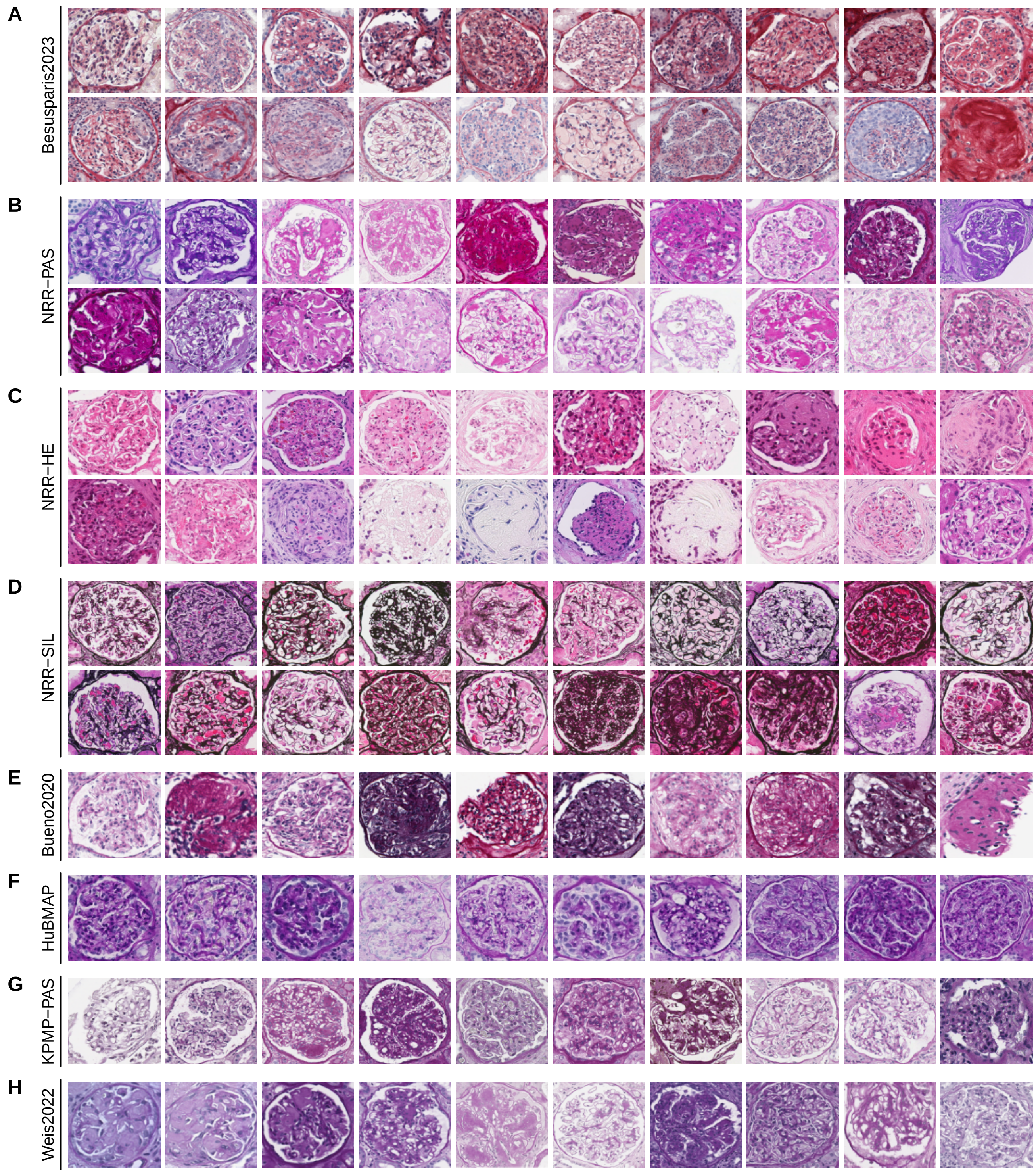

**Supp. Fig. 1.** Examples of glomerular image patches highlighting the stain variation within the eight datasets: Besusparis2023 (A), NRR-PAS (B), NRR-HE (C), NRR-SIL (D), Bueno2020 (E), HuBMAP (F), KPMP-PAS (G), Weis2022 (H). The 20 images presented in each of the panels A-D also served as the references used for stain-normalization of all datasets with the respective stain: mPSR (A), PAS (B), HE (C), and SIL (D).

#### Supplementary figure 2

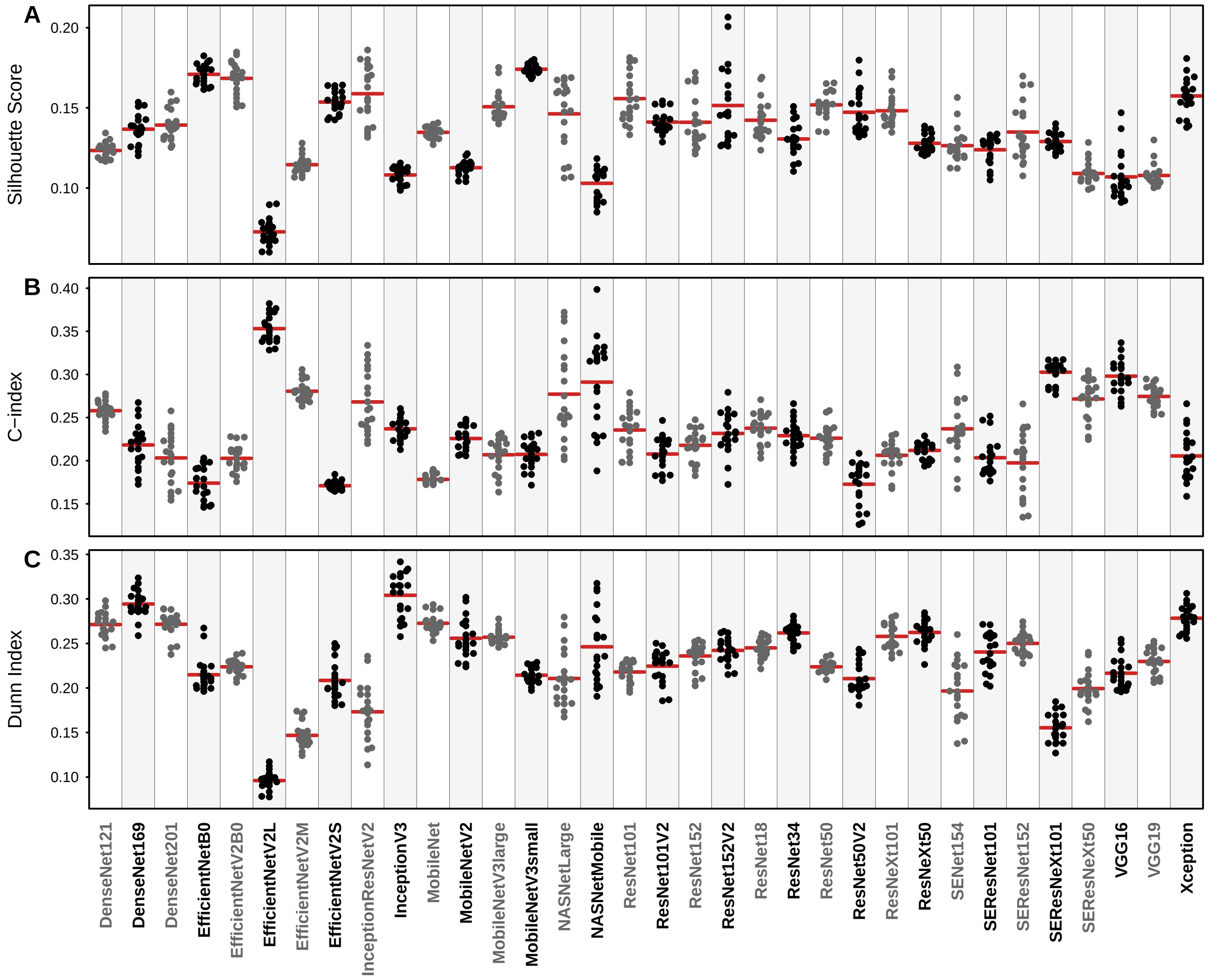

**Supp. Fig. 2.** Examples of the cluster validity index (CVI) results, evaluating the separation of GS and nonGS in the feature embedding of the KPMP-PAS images produced by different CNN models. The figure displays the results for three CVIs: the Silhouette Score (A), the C-index (B), and the Dunn index (C), where each data point for one of the indicated CNN models represents one of the 20 stain-normalized versions of the KPMP-PAS dataset. The red horizontal line for each dataset indicates the mean across the 20 respective scores.

#### Supplementary figure 3

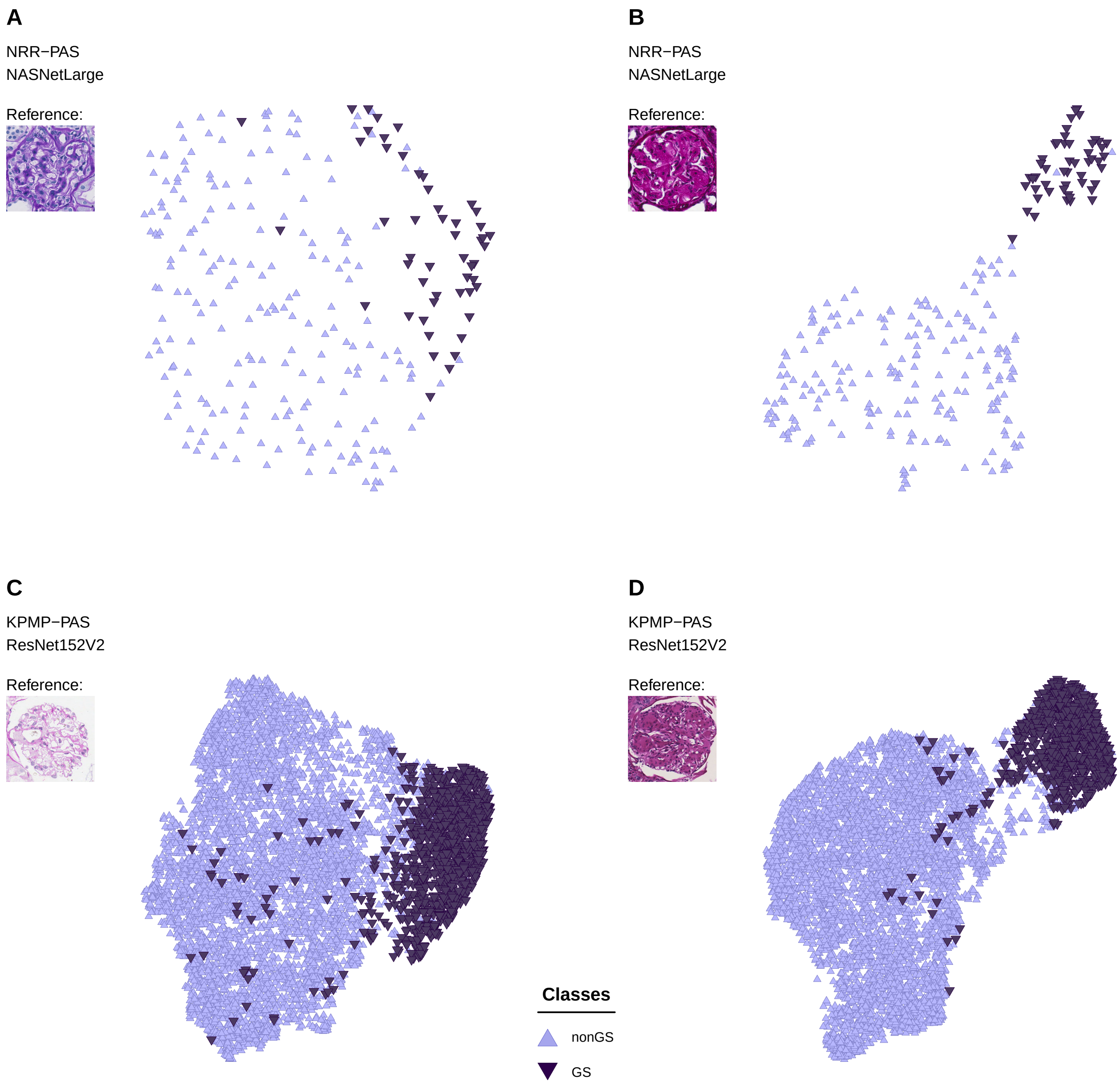

**Supp. Fig. 3.** Scatterplots illustrating differences in UMAP embeddings due to different stain-normalization references. The figure displays two different embeddings of both the NRR-PAS (A-B) and the KPMP-PAS (C-D) datasets, obtained by choosing two different reference images (shown as inlets in the corresponding plots) during stain-normalization of the respective dataset.

#### Supplementary figure 4

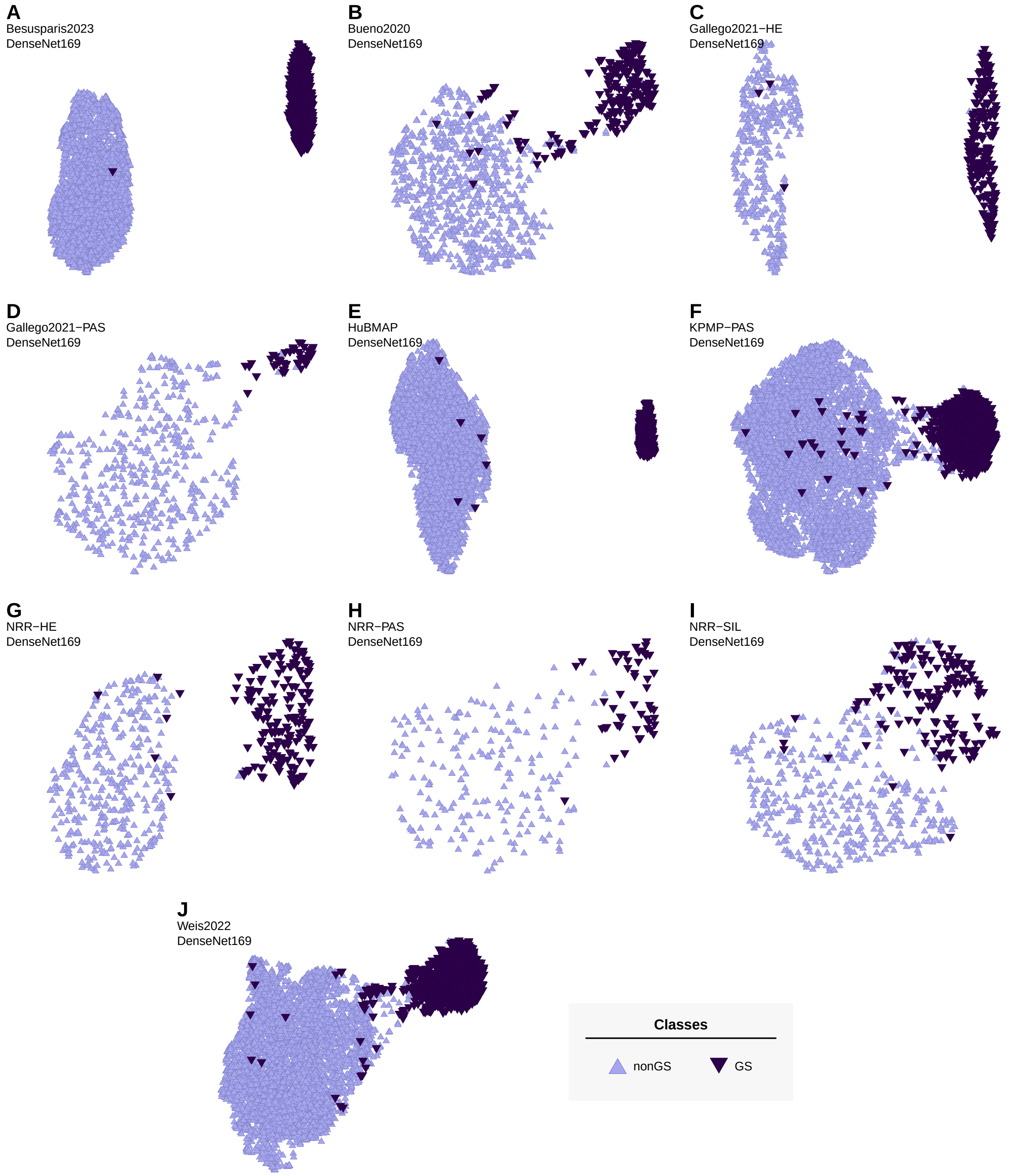

**Supp. Fig. 4.** Scatterplots illustrating the separation of GS and nonGS glomerular image patches following feature extraction with the *DenseNet169* and UMAP embedding in two dimensions. Each scatterplot depicts the feature embedding and classes of one stain-normalized version from one of the ten datasets: Besusparis2023 (A), Bueno2020 (B), Gallego2021-HE (C), Gallego2021-PAS (D), HuBMAP (E), KPMP-PAS (F), NRR-HE (G), NRR-PAS (H), NRR-SIL (I), and Weis2022 (J). Light and dark purple colors indicate nonGS and GS glomerular images, respectively.

#### Supplementary figure 5

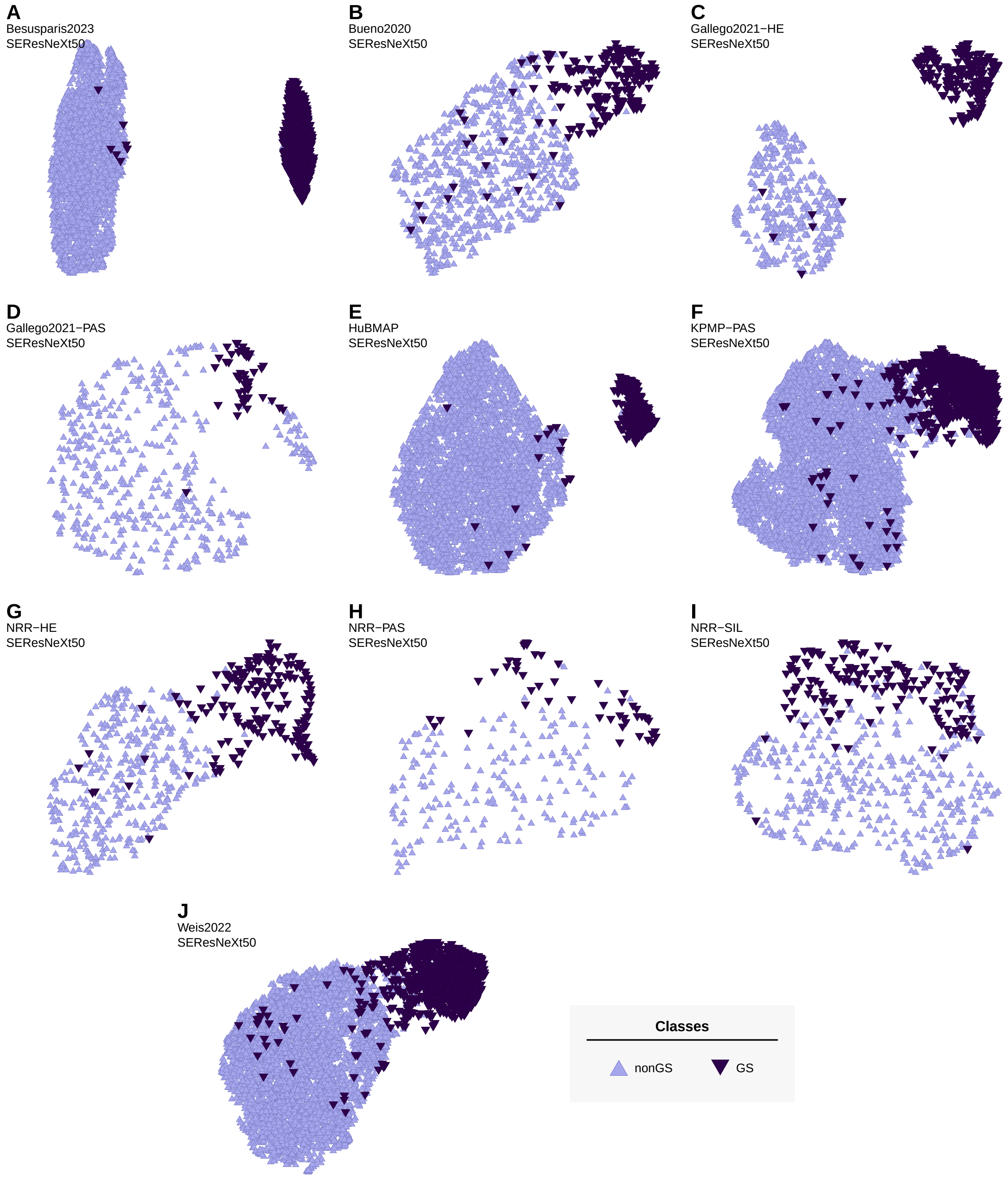

**Supp. Fig. 5.** Scatterplots illustrating the separation of GS and nonGS glomerular image patches following feature extraction with the *SE-ResNeXt50* and UMAP embedding in two dimensions. Each scatterplot depicts the feature embedding and classes of one stain-normalized version from one of the ten datasets: Besusparis2023 (A), Bueno2020 (B), Gallego2021-HE (C), Gallego2021-PAS (D), HuBMAP (E), KPMP-PAS (F), NRR-HE (G), NRR-PAS (H), NRR-SIL (I), and Weis2022 (J). Light and dark purple colors indicate nonGS and GS glomerular images, respectively.

#### Supplementary figure 6

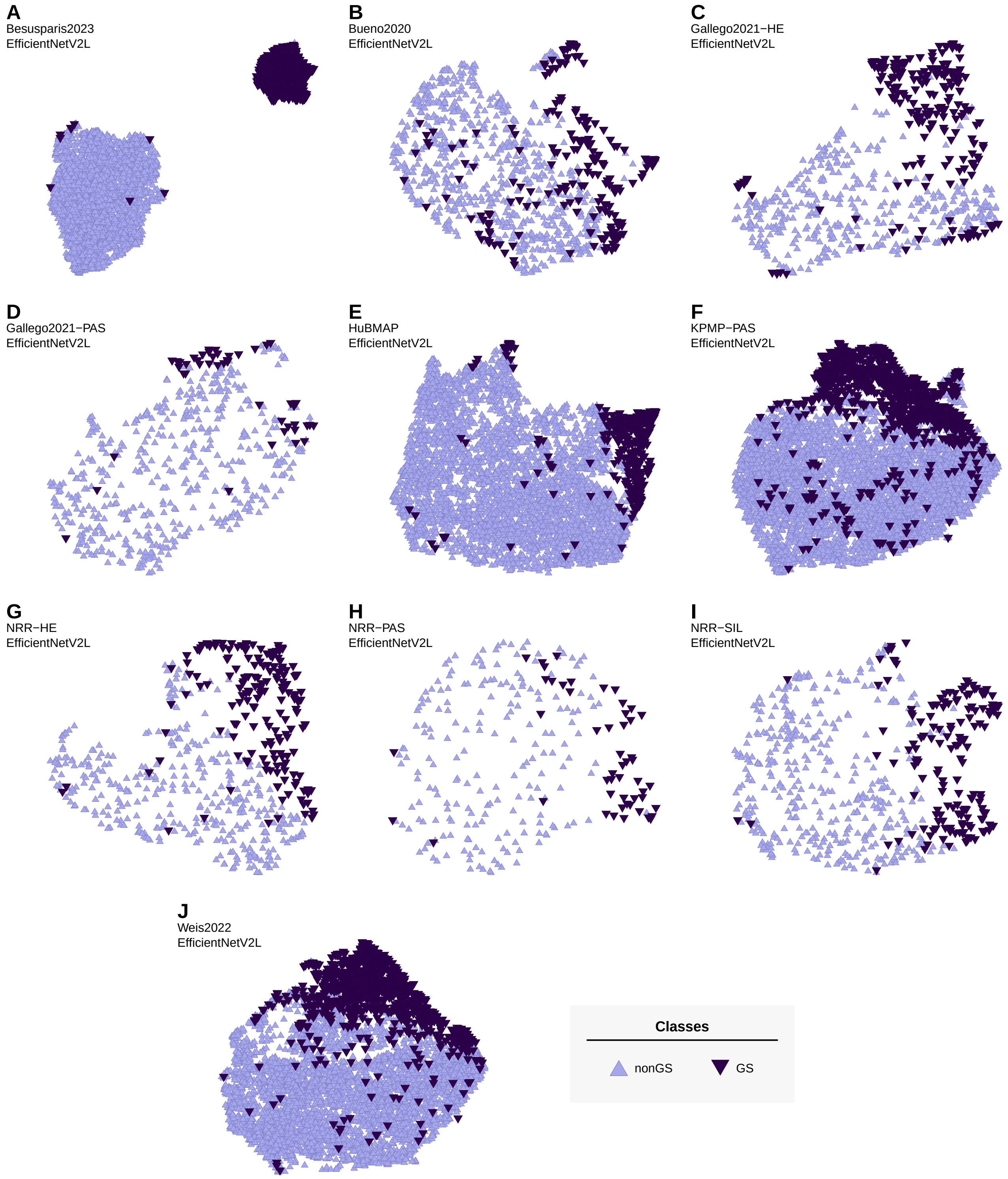

**Supp. Fig. 6.** Scatterplots illustrating the separation of GS and nonGS glomerular image patches following feature extraction with the *EfficientNetV2L* and UMAP embedding in two dimensions. Each scatterplot depicts the feature embedding and classes of one stain-normalized version from one of the ten datasets: Besusparis2023 (A), Bueno2020 (B), Gallego2021-HE (C), Gallego2021-PAS (D), HuBMAP (E), KPMP-PAS (F), NRR-HE (G), NRR-PAS (H), NRR-SIL (I), and Weis2022 (J). Light and dark purple colors indicate nonGS and GS glomerular images, respectively.

#### Supplementary figure 7

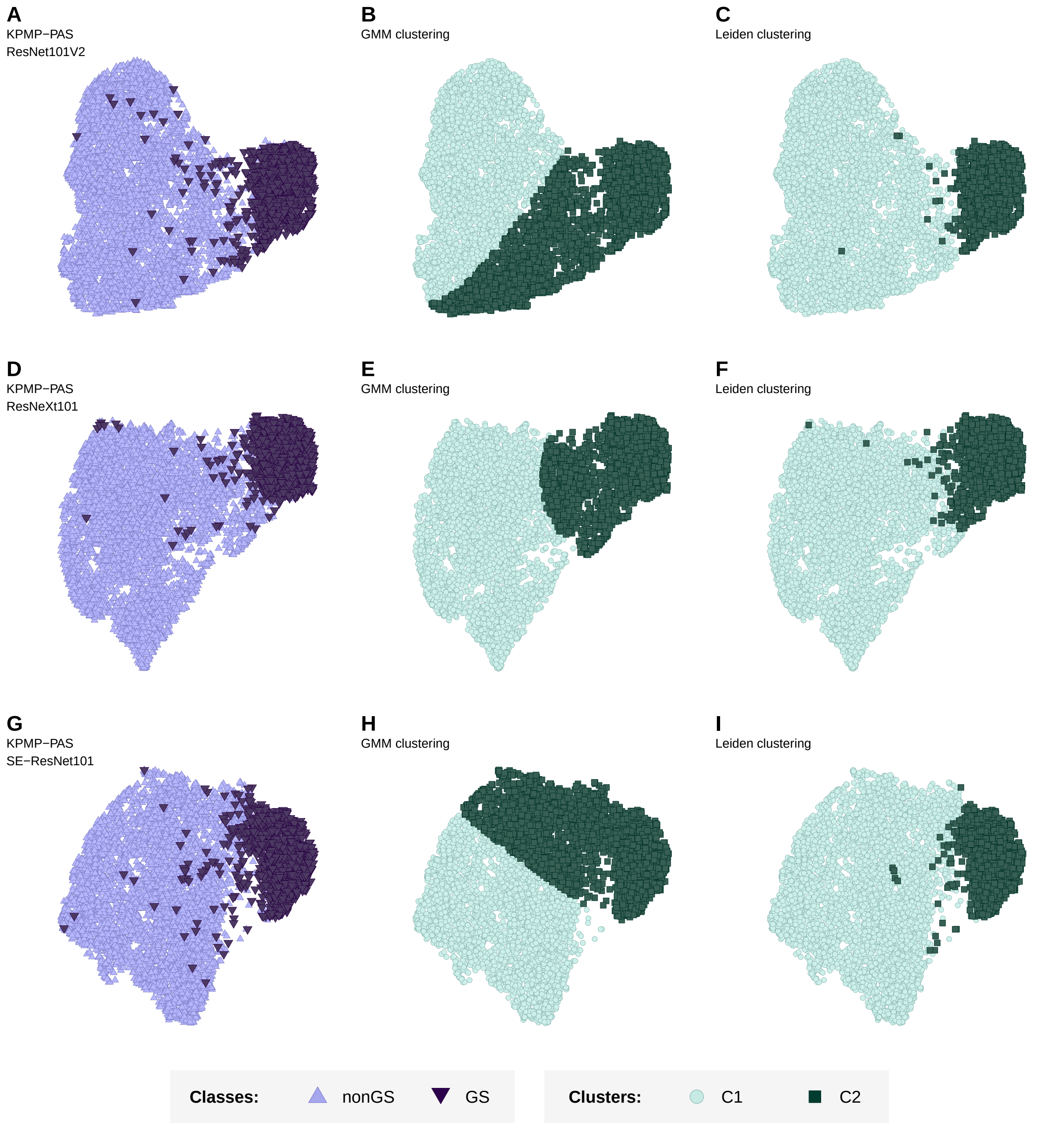

**Supp. Fig. 7.** Scatterplots visualizing some examples with pronounced differences in the clusters detected by the GMM and Leiden approaches. The figure displays the UMAP embedding of the KPMP-PAS images following the feature extraction with three different CNN models: ResNet101V2 (A-C), ResNeXt101 (D-F), and SE-ResNet101 (G-I). The leftmost column (A, D, G) illustrates the class affiliations in each UMAP embedding. The middle column (B, E, H) and right column (C, F, I) illustrate the cluster assignments produced by the GMM method and the Leiden method, respectively.

**Supplementary figure 8**

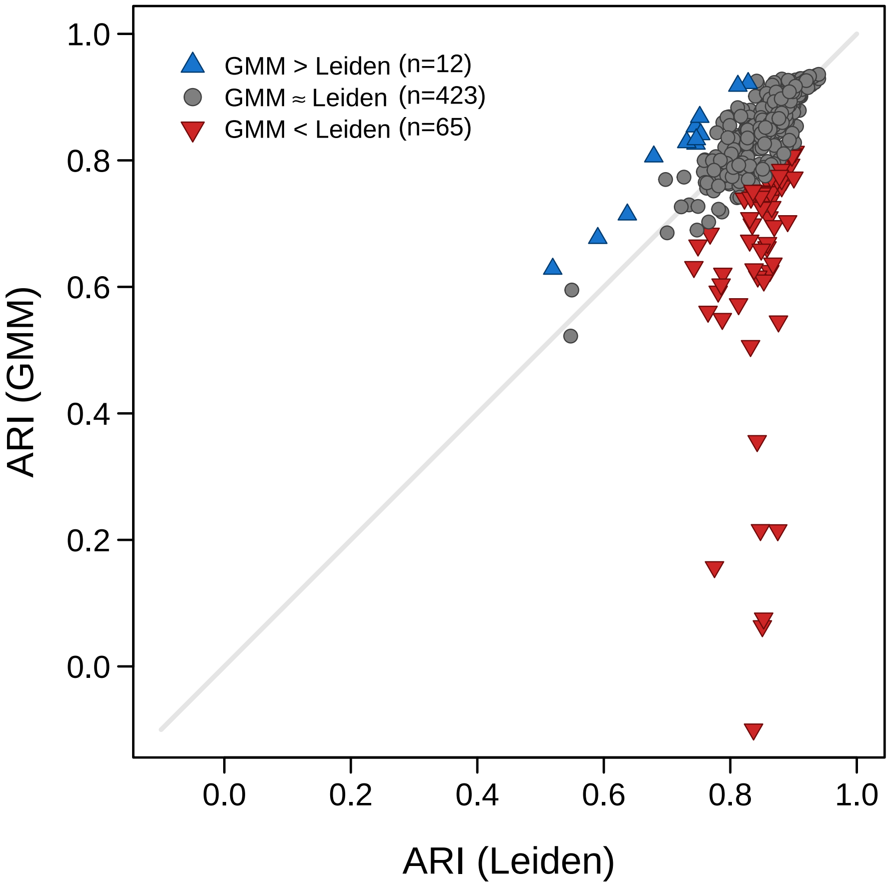

**Supp. Fig. 8.** Scatterplot comparing the ARI of GMM and Leiden clustering in accurately capturing the GS and nonGS classes. Each dot represents the results obtained for one combination of one of the 20 stain-normalized versions of the KPMP-PAS and one of the DenseNet, EfficientNetB0, EfficientNetV2B0, MobileNet, ResNet, ResNext, SE-ResNet, SE-ResNeXt, or SENet CNN models (excluding the other CNN models, which generally produced lower class separations in the feature embedding and thus also much lower quality clusterings regardless of clustering technique). Blue and red markers indicate cases in which the difference between GMM and Leiden is greater than 10% of the mean of both values, where blue indicates a better performance with GMM and red indicates a better performance with Leiden. Otherwise the performance was considered approximately equal (gray dots).

#### Supplementary figure 9

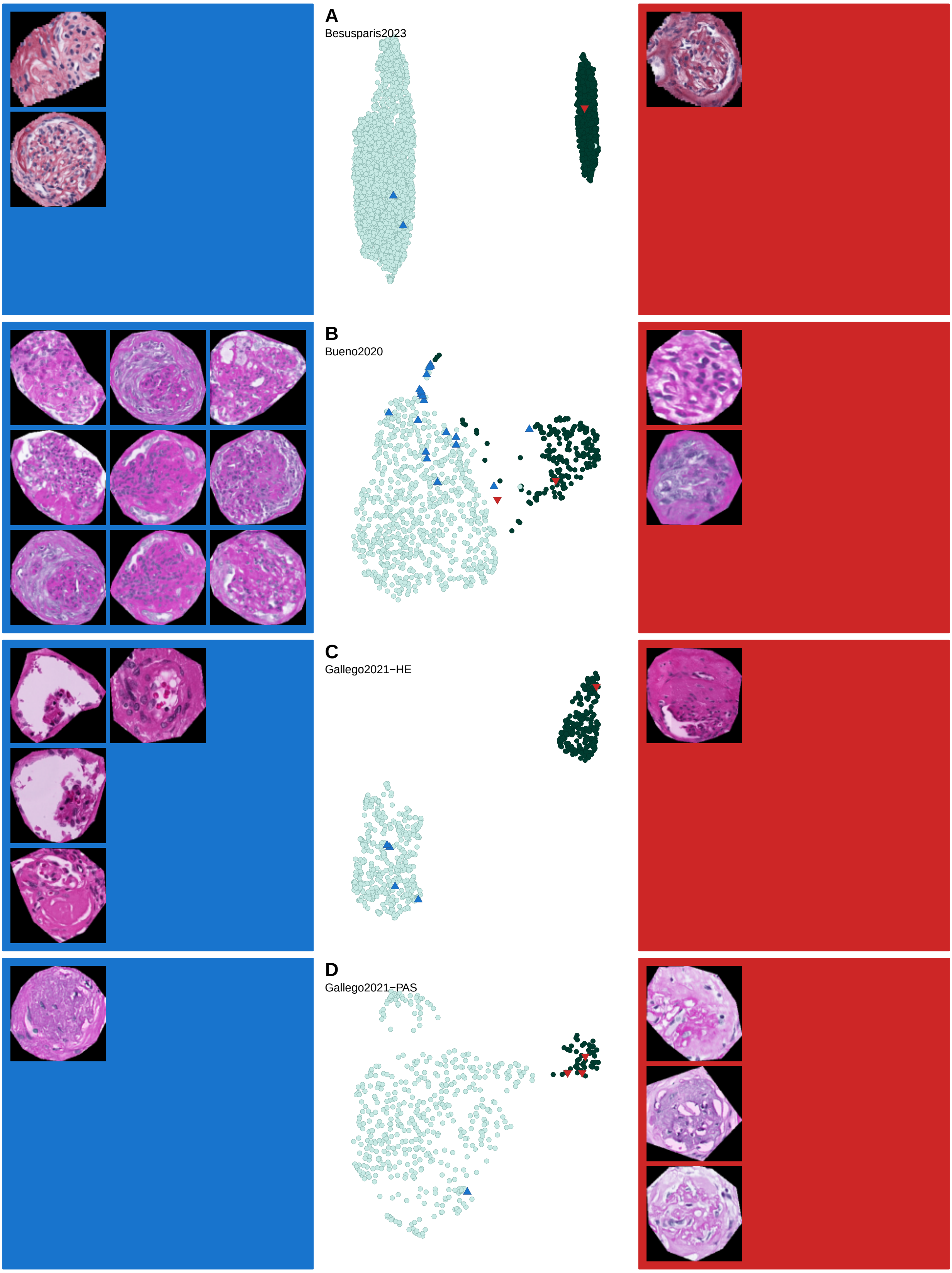

**Supp. Fig. 9.** Visual inspection of misclustered glomerular image patches in the four datasets: Besusparis2023 (A), Bueno2020 (B), Gallego2021-HE (C), Gallego2021-PAS (D). The UMAP embeddings of MobileNet-derived features are shown in the center column, with colors of individual data points indicating the clustering results: TNs (true negatives, light green), TPs (true positives, dark green), FNs (false negatives, blue), and FPs (false positives, red). The left and right columns illustrate up to nine examples of each of the FNs and FPs, respectively.

#### Supplementary figure 10

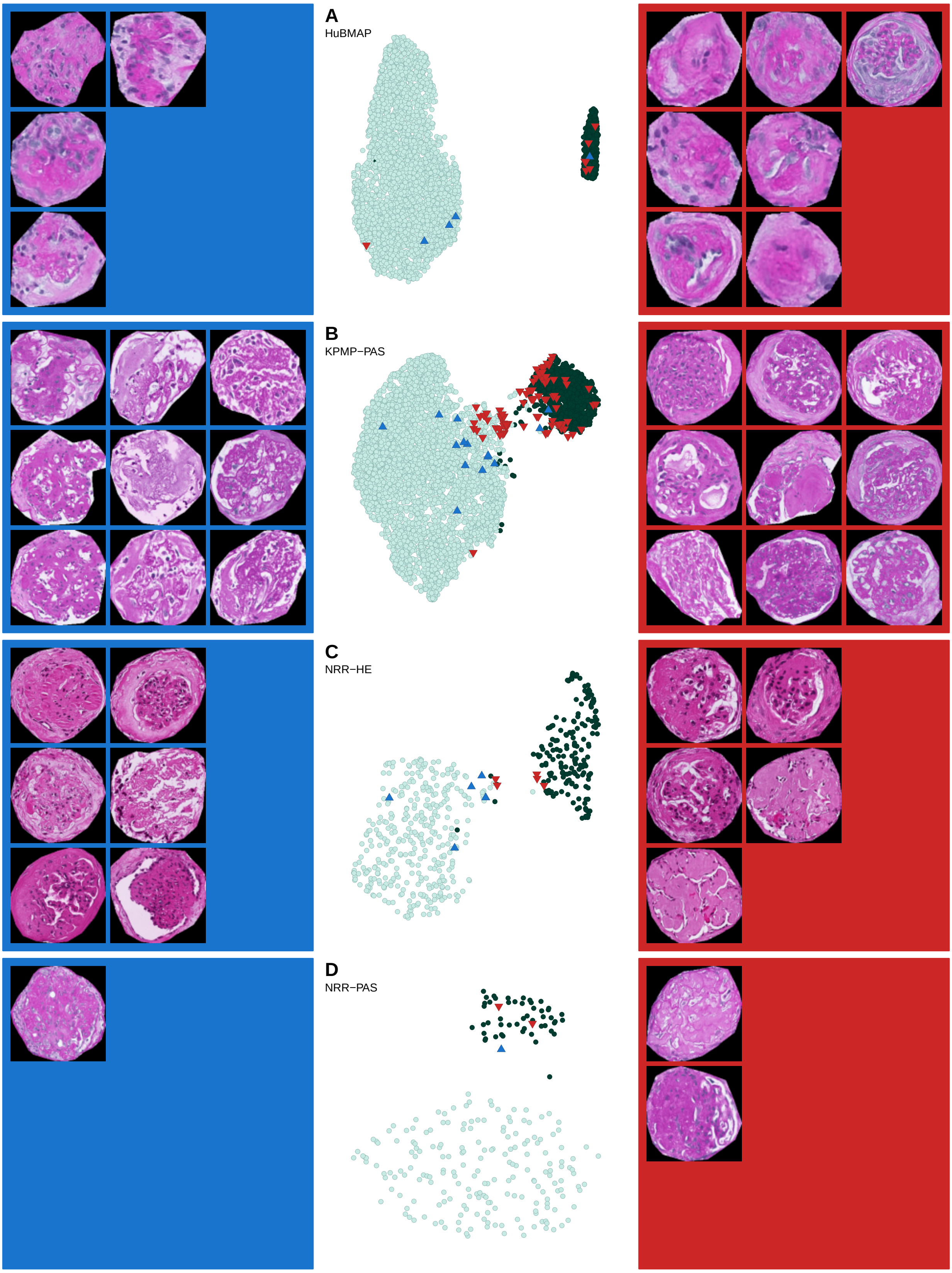

**Supp. Fig. 10.** Continuation of supplementary figure 9. Visual inspection of misclustered glomerular image patches in the four datasets: HuBMAP (A), KPMP-PAS (B), NRR-HE (C), NRR-PAS (D). The UMAP embeddings of MobileNet-derived features are shown in the center column, with colors of individual data points indicating the clustering results: TNs (light green), TPs (dark green), FNs (blue), and FPs (red). The left and right columns illustrate up to nine examples of each of the FNs and FPs, respectively.

#### Supplementary figure 11

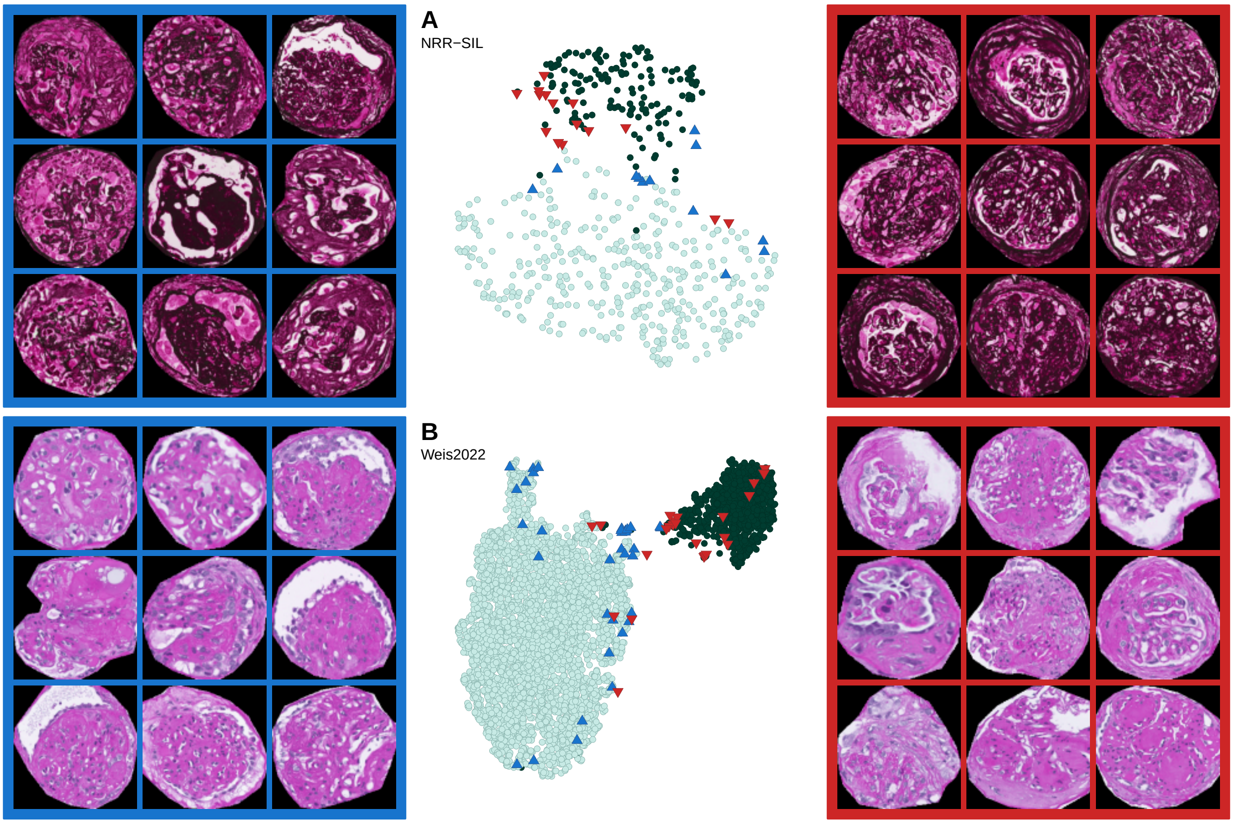

**Supp. Fig. 11.** Continuation of supplementary figure 9. Visual inspection of misclustered glomerular image patches in the two datasets: NRR-SIL (A) and Weis2022 (B). The UMAP embeddings of MobileNet-derived features are shown in the center column, with colors of individual data points indicating the clustering results: TNs (light green), TPs (dark green), FNs (blue), and FPs (red). The left and right columns illustrate up to nine examples of each of the FNs and FPs, respectively.

#### Supplementary figure 12

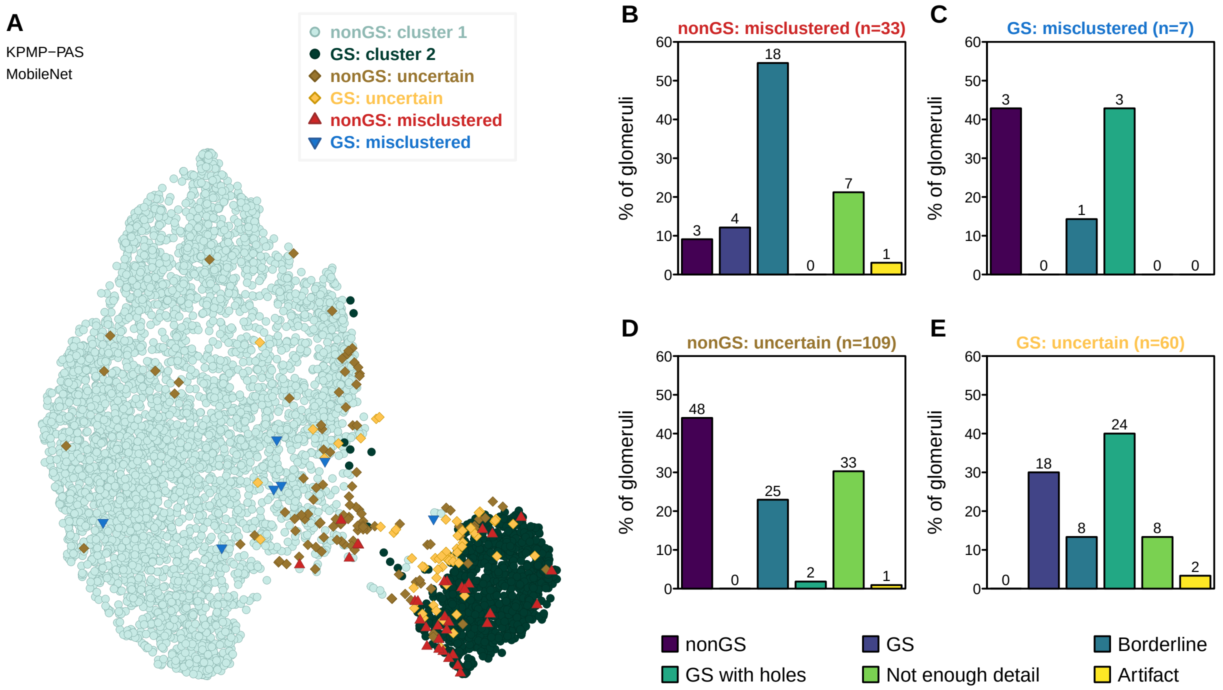

**Supp. Fig. 12.** Evaluation of incorrectly clustered glomerular image patches: voting over multiple CNN-derived feature embeddings. **A)** Consensus of the clustering of the KPMP-PAS dataset following feature extraction with three different CNN models (DenseNet169, MobileNet, and MobileNetV3small). Light and dark green circles indicate, respectively, glomeruli that are nonGS and always in cluster 1, and GS glomeruli that are always in cluster 2. Red and blue triangles indicate glomeruli that are always false-positives (nonGS in cluster 2) and false-negatives (GS in cluster 1), respectively. Brown and yellow diamonds indicate nonGS and GS glomeruli, respectively, that switch clusters between the different CNN models and can thus be identified as uncertain cases. **B-E)** Barplots representing a detailed analysis of the misclustered cases from each category, i.e. false-positives (B), false-negatives (C), and uncertain nonGS (D) and GS (E) glomeruli. Each barplot represents the percentages (absolute numbers above each bar) of glomeruli assigned by pathologists into the six categories: nonGS, GS, borderline glomeruli with advances sclerosis, GS with holes or split tissue, glomeruli with insufficient detail for labeling, and artifacts.
